# Identifying microbial species by single-molecule DNA optical mapping and resampling statistics

**DOI:** 10.1101/609412

**Authors:** Arno Bouwens, Jochem Deen, Raffaele Vitale, Laurens D’Huys, Vince Goyvaerts, Adrien Descloux, Doortje Borrenberghs, Kristin Grussmayer, Tomas Lukes, Rafael Camacho, Jia Su, Cyril Ruckebusch, Theo Lasser, Dimitri Van De Ville, Johan Hofkens, Aleksandra Radenovic, Kris Pieter Frans Janssen

## Abstract

Single molecule DNA mapping has the potential to serve as a powerful complement to high-throughput sequencing in metagenomic analysis. Offering longer read lengths and forgoing the need for complex library preparation and amplification, mapping stands to provide an unbiased view into the composition of complex viromes and/or microbiomes. To fully enable mapping-based metagenomics, sensitivity and specificity of DNA map analysis and identification need to be improved. Using detailed simulations and experimental data, we first demonstrate how fluorescence imaging of surface stretched, sequence specifically labeled DNA fragments can yield highly sensitive identification of targets. Secondly, a new analysis technique is introduced to increase specificity of the analysis, allowing even closely related species to be resolved. Thirdly, we show how an increase in resolution improves sensitivity. Finally, we demonstrate that these methods are capable of identifying species with long genomes such as bacteria with high sensitivity.

## INTRODUCTION

Communities of microbial species and their collective genes, typically referred to as a microbiome, have become an increasingly important study area over the last years. The interplay between highly diverse microbial species and their hosts turned out to play a key role in many fields of biology. (1) So not surprisingly, recent years have seen a tremendous interest in the microbiome, especially in the gastrointestinal tract and its relatedness to various diseases. (2–4) The microbiome does not only encompass bacterial species, it also covers fungal and viral communities. It is reported that human feces contain at least 10^9^ virus-like particles (VLPs) per gram, a majority of which are bacteriophages. (5) The effect they have on the microbiome can be significant: as an example, a recent article showed how type I diabetes was associated with a significant reduction in the abundance of *Circoviridae*-related sequences. (6)

The classical method to identify microbial species is by culturing them in laboratory conditions. (7) However, this method is extremely biased since the overwhelming majority of species is currently unculturable. (8) Newer approaches for identification have focused on identification by sequencing, the most common of which targets a universally conserved marker gene on the genome (like the 16s ribosomal DNA) and amplifies it. Still, identification by amplification suffers from some important drawbacks. First, amplification tends to introduce biases in the sample, skewing abundance measurements. (9) Second, it only groups together species that share the same amplification gene. And finally, it leaves viruses *invisible* because of the absence of a universally conserved marker gene in their genome. (10) A second approach is to sequence the whole genome of all the species in the microbiome. (9) While whole-genome sequencing does not suffer from the same problems such as amplification bias and intrinsic preclusion of viruses, the most common sequencing techniques are unable to deliver both long sequence-reads and high-throughput. (10) Short contigs (contiguous sequences), for example, make identification difficult because of missing long-range structural information.

As a promising alternative, DNA mapping can be used for the same purpose. (11) Instead of providing sequence data, DNA mapping provides long-range location density of specific short sequences on the genome. Mapping strategies have been used to match DNA maps to, for instance, plasmids for bacterial resistance studies (12), to bacteriophage sequences (13), *Escherichia Coli* (14), and to CRISPR-CAS9 edited regions in bacterial genomes. (15) More recently, DNA mapping was used to identify genomic regions in individual cells. (16) The most common approach for DNA mapping is to enzymatically attach fluorescent labels to specific short sequences on the DNA using DNA methyltransferases (17–19) and image them by a fluorescence microscope. Other labeling methods transfer labels using DNA nickases, (20) or intercalating dyes using either competitive binding with an inhibitor (21) or denaturation-renaturation mapping. (22)

In order to identify a species using DNA mapping, the measured DNA maps need to be compared or *matched* to the expected DNA maps for that species, which is constructed from the species’ known sequence. The current matching approaches can be roughly categorized in two classes: (1) methodologies that rely on cross-correlating the measured profile with an expected profile and (2) methodologies that use dynamic-programming-based algorithms that compare the distance between labels (in basepairs) in the measured and a theoretical map. (23–25) The former handles the DNA map as an intensity profile over the DNA strand, while the latter handles the DNA map as the list of the labelled positions on the DNA strand. Both types of methods return a so-called matching score that increases when the measured map more closely resembles the expected or theoretical map.

An essential step in species identification by DNA mapping is deciding whether the matching score is high enough to assign the measured DNA map to a genome or species. One approach, based on dynamic programming, uses a quality index for rejecting false assignations. (26) This quality index is based on the ratio between the matching score and the standard deviation of false matching scores at other positions in the genome. In a second approach, a *p*-value is calculated for the probability that the matching score is randomly generated. (27) If this probability is low (e.g., below 5%), the matching score is considered to be statistically significant and therefore reliable. Recently, Nilsson *et al*. suggested the use of a *p*-value combined with an information score to determine the quality of a match. Their *p*-value is calculated by imposing a *hard null-*model on the distribution of the matching score (14), which makes it prone to error if the model conditions are not satisfied. Moreover, *p*-values calculated for matches to different species cannot be compared to each other. It is therefore not clear how to discriminate between closely related species when a single DNA map matches significantly to multiple species.

In this article, we propose a new technique rooted in resampling statistics for assigning measured double-stranded DNA maps to microbial species. Our technique is applicable to either cross-correlation and dynamic programming matching methods. We investigate the performance of DNA mapping by validating our method on bacteriophage identification and show how it can be generalized to bacterial species. To compare the performance of different imaging approaches for DNA mapping, we performed a series of experiments, and developed a simulation tool that closely mimics the sample preparation as well as wide-field and super-resolution microscopy techniques.

## MATERIALS AND METHODS

### M.TaqI directed labeling using a Rhodamine B-tagged SAM analogue

DNA from bacteriophage lambda (Thermo Scientific) and T7 (Yorkshire Bioscience) and from bacterium *Vibrio Harveyi* (*V. Harveyi*, ATCC) was enzymatically labelled at a final concentration of 50 ng/µl, using 35 µM of the rhodamine B functionalized AdoMet analogue and 0.14 mg/ml M.TaqI methyltransferase enzyme (recognition sequence 5’-TCGA-3’). The reaction was carried out at 60°C for 2 hours in a custom labeling buffer with a final concentration of 50 mM K-acetate (Sigma), 10 mM Mg-Acetate (Sigma), 20 mM MES (Sigma) and 0.1 mg/ml BSA (Sigma), buffered at pH 5.75. Subsequently, 2 µl of proteinase k (800 units ml-1, NEB) were added and reacted for 1 hour at 50°C. Finally, the reaction product was purified using CHROMA SPIN+TE-1000 columns (Clontech, Takara Bio).

### Preparation of Zeonex coated coverslips

22×22×0.17 mm #1.5 glass coverslips (Menzel-Gläser) were rinsed thoroughly with water and blown dry. Next, the coverslips were thermally treated overnight at 450 °C and subjected to a 30 minutes UV-ozone treatment in a UVP PR-100 ozone cleaner (UVP, Upland, California, USA). A small volume of 1.5 % w/v Zeonex (Zeon Chemicals L.P.) in Toluene (Acros Organics) was subsequently deposited using a spin coater, programmed to rotate 15 seconds at 1000 RPM, 1 minute at 10000 RPM and 10 seconds at 1000 RPM. The Zeonex solution was always sonicated for 40 minutes prior to coating. Finally, coated coverslips were dried at 110 °C for 2 hours prior to storage in a desiccator at room temperature.

### Stretching of labelled DNA molecules on Zeonex coated coverslips

Purified DNA was dissolved in 50 mM MES (pH 5.6), and deposited in stretched conformation by mechanically dragging a 2 µl droplet over the surface of a Zeonex-coated coverslip at a speed of 4.4 mm/min using a disposable pipet tip, as described earlier. (28) Stretched samples were stored dry and were vacuum dried overnight prior to imaging.

### Imaging

Imaging was performed with a Zeiss SIM Elyra microscope with a Zeiss Plan-APOCHROMAT 63x oil immersion objective (numerical aperture 1.4) and an EMCCD camera (exposure time 300 ms/frame, EM gain setting 35). An extra 1.6x image magnification was applied. The field of view per image was 75 × 75 µm^2^. The camera pixel size projected in the sample was 80 nm/pixel. The 561 nm excitation laser provided a power of ∼ 3mW over the field of view. Fluorescence emission was filtered by a 570-620 nm band-pass filter. For each field of view, 25 frames were recorded for 5 SIM modulation angles and 5 phases/angle. The illumination patterns for SR-SIM were created by a grating with a period of 34 µm. A drop of milliQ water was placed on top of the sample before imaging (see Supplementary Information, Section 2.5). A wide-field image was calculated by averaging over the 25 frames. SR-SIM reconstruction was done with the open-source fairSIM plugin for ImageJ. (29) DNA fragments were segmented manually on the SR-SIM images using ImageJ. For each imaged DNA fragment, both wide-field and SR-SIM signals were extracted.

### Calculation of the matching score

For each species of interest, the cross-correlation function *XC*(*δ, f, d*) between the measured DNA map and the expected DNA map was calculated. The cross-correlation function is a measure of the similarity between two signals as a function of the displacement of one with respect to the other. As the measured DNA map is overstretched during the experimental procedure and can have both 5’-3’ and 3’-5’ orientations, the cross-correlation function is calculated varying the overstretching factor of the expected DNA map, *f*, between 1.7 and 1.76 with steps of 0.01 and the orientation of the measured DNA map, *d*, as:

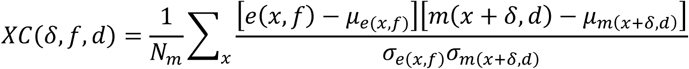

where *e*(*x, f*) and *m*(*x* + *δ, d*) represent the expected and the measured DNA map, respectively, *δ* quantifies the displacement between the two, *μ*and *σ* denote mean and standard deviation, and *N*_*m*,_ corresponds to the total number of sampling points in *m*(*x* + *δ, d*). The expected DNA maps were constructed from the known DNA sequences of the tested microbial species (downloaded from the NCBI database). Specifically, the *n* locations of each methyltransferase enzyme recognition sequence (in units of bp) were listed. This list, *l*(*n*), was converted into the intensity signal *e*(*x, f*) by summing up Point Spread Functions (PSFs) centered at each recognition sequence location. The signal was sampled along the DNA with a step size equal to the camera pixel size projected in the sample:

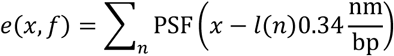

where PSF(*x*) is the microscope PSF and *p* the projected pixel size. The final matching score *S* was taken as the global maximum across all *δ* and *f* values, and for both orientations *d*:

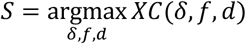

This way, the optimal shift, orientation and overstretching factor of the measured DNA map were also found.

### Significance test on the matching score (“matching significance test”)

To test the statistical significance of a matching score, we applied permutation testing to calculate an empirical *p*-value. Per each species, we constructed randomized DNA maps starting from the expected DNA map and randomly reshuffling the locations of the labels. A new matching score was calculated for each reshuffled DNA map. This reshuffling is carried out a large number of times (say *N*_*p*_ = 10^4^). Finally, a *p*-value, *p*_1_, was calculated as:

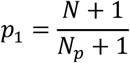

where *N* is the number of times the randomized matching score was found to be higher than the measured matching score. In this way, the tested matching score is contrasted against the distribution of randomized matching scores. If the tested matching score is found to be systematically higher than those resulting from such a randomization (i.e. *p*_1_ is found to be lower than a certain significance threshold, *α*_1_), the match is considered statistically significant and retained as such. The choice of the significance threshold allows one to trade identification sensitivity against identification specificity: a low threshold results in lower sensitivity and higher specificity and vice versa.

For the sake of clarity, the fluorescent label reshuffling was carried out in a windowed manner. Instead of randomizing the label locations over the full length of the reference genome, we subdivided it into 10 kb-long windows and the labels were reshuffled within these intervals. This way, some of the larger-scale DNA structures were preserved, which improved the assignation specificity while having just a little effect on the assignation sensitivity.

### Resampling the highest matching score to improve specificity (“resampling step”)

To compare matching scores for different species and reject those that are significantly lower than the highest significant one, we resampled its corresponding expected DNA map as follows. From the list of its label locations, we randomly removed 2 labels, creating a resampled DNA map. Labels were removed only in the region of the expected DNA map where the measured DNA map was found to match after the matching significance test. A resampled matching score was found by calculating the matching score between the measured DNA map and the resampled DNA map, without re-optimizing the shift, orientation and overstretching factor found in the previous step. This resampling procedure was repeated enough times (say *N*_*r*_ = 4000) to create a well-sampled estimate of the spread on the highest significant matching score.

For any other matching score that was found significant after the matching significance test, a second *p*-value, *p*_2_, was calculated:

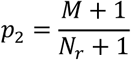

where *M* is the number of times the resampled score was found to be lower than the tested matching score. When *p*_2_ is lower than an imposed significance threshold, *α*_2_, the tested matching score is considered significantly lower than the highest significant matching score. In this case, the respective match can be discarded as it is significantly worse than the best observed match. On the other hand, when a matching score is not significantly lower than the highest significant matching score, it is retained and the corresponding DNA map assigned to more than one species.

### Simulating DNA optical mapping

We developed a simulation model, implemented in MATLAB, covering the entire process of DNA mapping, including strand breakage, DNA labeling, DNA deposition and imaging.

First, the DNA sequences of bacteriophage species were downloaded from the NCBI database. Next, DNA fragments of 35 kbp length were drawn from the full DNA sequences, with random starting position. This step simulates the random breakage of the DNA which occurs due to shearing forces from pipetting and the presence of nucleases in the sample mixture. The length of 35 kbp corresponds to a typical DNA fragment length in our experiments (see Supplementary Information, Section 2.2). 1000 DNA fragments were simulated per species.

In earlier work we found that the methylation enzyme does not label all recognition sites on the DNA, but instead has a labeling efficiency of around 80%. (30) Moreover, we noticed that this efficiency can be lower when the synthetic cofactor has degraded prior to labeling or when there are remnants of the natural cofactor AdoMet present in the labeling mixture. In this study, we set the simulated labeling efficiency to 75%. Furthermore, the enzyme sometimes transfers a label to an incorrect sequence, resulting in a false positive label. Here, labeling efficiency and false positive labeling were simulated as Poisson processes, yielding a specific list of simulated label positions on each DNA fragment. Linearization of the DNA fragments was simulated by applying a constant overstretching factor of 1.75 to the list of simulated label positions. This overstretching factor corresponds to the value found in experiments. (28)

Finally, imaging of the DNA fragments was simulated. The PSF of the microscope was approximated by a 2D Gaussian with Full-Width at Half-Maximum (FWHM) of 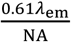, where NA is the microscope numerical aperture (1.4 in our experiments), and *λ*_em_ the fluorescence emission wavelength (576 nm for the rhodamine-B dye used here). The simulated images were sampled according to the microscope pixel size. Photon emission from the fluorescent labels was modeled as a Poisson process. The average number of collected photons per fluorescent label and per camera exposure time (300 ms) was estimated to be ∼10^4^. For SR-SIM, 25 frames were simulated with sinusoidal illumination patterns with a modulation depth of 0.7 (corresponding to the value observed experimentally), and having the same 5 orientations and 5 phases/orientation as in the experiments. The EMCCD camera quantum gain, thermal noise, and read-out noise were simulated and calibrated following Reference (31). For simulated wide-field microscopy, the 25 frames were averaged. As for the experiments, SR-SIM reconstruction was done with the open-source fairSIM plugin for ImageJ. (29)

## RESULTS AND DISCUSSION

To assess and validate our new method for identifying species by optical DNA mapping, we used three complementary test case-studies, two consisting of experimental data measured on bacteriophage and bacterial DNA, respectively, and one consisting of simulated data from a detailed model. The simulated data allowed us to find out what parameters are important when identifying species based on DNA maps.

The experimental data were obtained by imaging DNA fragments from bacteriophages lambda and T7 and bacterium *V. Harveyi*. We labeled DNA fragments with fluorescent dyes using the M.TaqI methyltransferase enzyme with recognition sequence 5’-TCGA-3’ and a synthetic AdoMet cofactor. (17, 32) After labeling (Figure 1A), the DNA was linearized on a coated coverslip using a “rolling droplet” technique (Figure 1B). This technique causes the DNA fragments to be overstretched by a factor of 1.7 to 1.75. (28) We imaged the labeled DNA with a wide-field microscope. Finally, we extracted the fluorescence intensity signal along the linearized DNA for each individual DNA fragment (Figure 1C).

**Figure 1:**
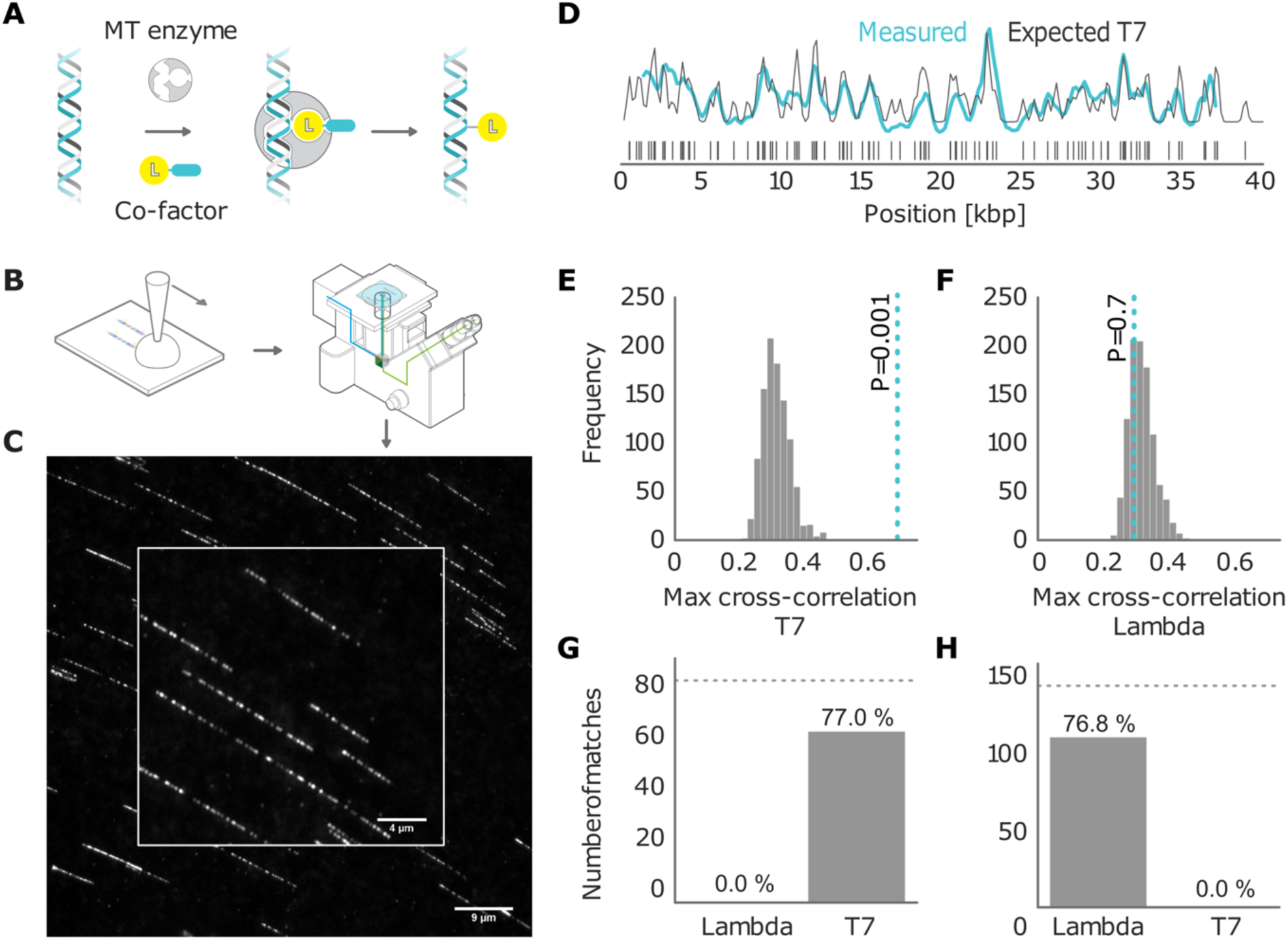
(A) Graphical sketch of the enzymatic labeling procedure. (B) After enzymatic labeling, DNA fragments are surface deposited and overstretched using a “rolling droplet” procedure, (28) followed by fluorescence imaging. (C) Representative image of labeled T7 DNA molecules stretched on a coated coverslip, obtained by wide-field fluorescence microscopy imaging. (D) Measured DNA map of one of the imaged molecules (cyan) overlaid with the T7 expected DNA map (black). (E, F) Histogram of the randomized matching scores corresponding to the maximum cross-correlations of the measured DNA maps with the reshuffled DNA maps. The red line indicates the matching score found from the maximum cross-correlation of the measured DNA map with the expected DNA map, with low *p*_1_-value for T7 (ground truth) (E) and high *p*_1_-value for lambda (control) (F). (G) Results of matching 87 T7 DNA molecules imaged by wide-field microscopy to the expected DNA maps of bacteriophages lambda and T7 (*α*_1_ = 0.001). (H) Results of matching 142 lambda DNA molecules imaged by wide-field microscopy to the expected DNA maps of bacteriophages lambda and T7 (*α*_1_ = 0.001). The dotted line indicates the total amount of DNA maps concerned.

The simulated data allowed us to investigate the performance of our identification method on expanded sets of species. As outlined before, the simulation model generates DNA maps for these species accounting for the various sources of experimental variation: enzymatic labeling of the DNA, overstretching and shearing of the DNA, fluorescence photo-physics, and imaging.

### Assigning optical DNA maps to bacteriophages

In this section, the main objective of our method will be to correctly assign every measured DNA map to a bacteriophage from a set of candidate species. To this end, we calculated the cross-correlation of the measured fluorescence signal with each of the expected signals for the candidate species. We considered the maximum of the cross-correlation function as the matching score. The expected intensity signals were calculated by summing up the microscope PSF centered at each of the locations of the labeling enzyme recognition sequence, given the full genome of the bacteriophages of interest (Figure 1D).

For every measured DNA map, we calculated the matching scores for all the candidate species. In order to determine which of these scores should be considered as reliable matches, we subjected all matching scores to a significance test. The goal of the test is to reject matches where the score is not significantly higher than the scores for randomized DNA maps, which are generated by reshuffling the dye locations on the expected DNA map. As described in the Materials and Methods section, we used permutation testing to calculate a corresponding *p*-value (*p*_1_). If *p*_1_ is found to be lower than a certain significance threshold, the matching score is retained as significant (Figure 1E). If not, the matching is rejected (Figure 1F). The result of this procedure is a list of species that yield a significant match for the measured DNA map.

Figures 1G and 1H show the results of this test for experimental data recorded with wide-field microscopy of overstretched DNA fragments from bacteriophage T7 (87 measured maps) and lambda (142 measured maps). The DNA fragments were assigned to the right bacteriophage species: the fraction of DNA fragments that were assigned to the ground truth species (true positives) is above 75%, while false positives are at 0%. The significance threshold was set at *α*_1_ = 0.001.

While the goal of this significance test is the same as in the method of Nilsson *et al*. (14), our approach is non-parametric as we do not need to assume any statistical distribution for the matching score in order to calculate *p*_1_. Hence, any bias due to assumption inaccuracy is inherently avoided. Furthermore, our method can easily be applied to other types of matching scores, for example, the alignment score from dynamic programming methods. (23) This matching significance test is therefore a widely-applicable matching reliability metric. In addition, testing the statistical significance of the matching score allows the whole approach to be robust against biased definitions of the database of reference species. That is to say that if fragments drawn from a species which is not included in the reference database are imaged, we expect our methodology to recognize them as unknowns (i.e. no significant matchings should be returned for any of the reference species under assessment). This is a clear advantage over just assigning these fragments to the species for which a maximum matching score is found as it reduces the impact of false positives on the final outcomes. Similarly, in situations in which DNA maps which are shared by multiple microbial species (due e.g. to a high evolutionary similarity) are imaged, they will not be necessarily assigned to only one of them, allowing common genomic subsequences to be therefore easily recognised

### Improving sensitivity by overstretching and super-resolution microscopy

For a labeling enzyme with a 4-base recognition sequence, the expected number of labels in a random string of nucleotides is 1 per 256 bases, or about 1 label every 87 nm of full-length DNA. Therefore, one can expect to have more than 1 fluorescent dye in a diffraction-limited PSF spot in about 93% of the cases^i^. Signal overlap from different labels limits the amount of information that can be extracted from the DNA sequence. We can therefore expect improved identification of bacteriophages when the PSF is narrowed. Besides improving the optical resolution, another way in which the effective width of the PSF can be narrowed (in terms of basepairs) is by overstretching DNA. This approach is equivalent to expansion microscopy. When depositing its molecules on a surface using DNA combing, the DNA adopts an overstretched configuration where the basepair distance increases from 0.34 nm to 0.6 nm. This overstretching of around 75% corresponds to the maximal length of double-stranded DNA before strand-breakage. (28, 33) Consequently, overstretching reduces the fluorescent dye overlap probability to 81%. Super-resolved microscopy techniques can further narrow the PSF, thus reducing the overlap probability to 54%.

To investigate the effects of improved resolution on bacteriophage identification, we simulated four different scenarios: (1) diffraction-limited wide-field microscopy of unstretched DNA; (2) wide-field microscopy of overstretched DNA (stretching factor 1.75); (3) Super-Resolved Structured Illumination Microscopy (SR-SIM) of overstretched DNA; (4) Single Molecule Localization Microscopy (SMLM) with reverse photobleaching of overstretched DNA. These methods progressively increase the effective resolution. SR-SIM is capable of improving the resolution about two-fold beyond the diffraction limit. (34) In SMLM with reverse photobleaching, all the dyes on the DNA are localized and fitted with a 2D Gaussian with a FWHM set to the accuracy of the localization resulting from a super-resolution image. (19, 35) The accuracy of localization using this method was 20 nm (see Supplementary Information, Section 3.2). When generating the simulated DNA maps, we also included the imperfections of the labeling procedure.

We generated 1000 DNA fragments for each bacteriophage from a set of 10 species, of which 6 are from the same family as bacteriophage lambda (Siphoviridae), 2 from the same family as bacteriophage T7 (Podoviridae), and 2 from the Myoviridae family. Because different species can contain widely different labeling densities we included two bacteriophages with high labeling density (17.9 and 10.9 sites per kbp) and two with low labeling density (1.0 and 1.1 sites per kbp). The average labeling density of the other species is ∼3 sites per kbp. For a detailed overview of the selected species see Supplementary Table 1.

The simulated DNA molecules were matched to all of the 10 bacteriophage species, and *p*_1_-values were calculated for the matching scores. If a match passed the significance test with threshold *α*_1_ = 0.001, the DNA molecule is assigned to the corresponding species of bacteriophage. In this way, an assignation matrix of significant matches was constructed, as shown in Figure 2B. A perfect assignation matrix would show 100% matches on the diagonal and 0% elsewhere.

**Figure 2:**
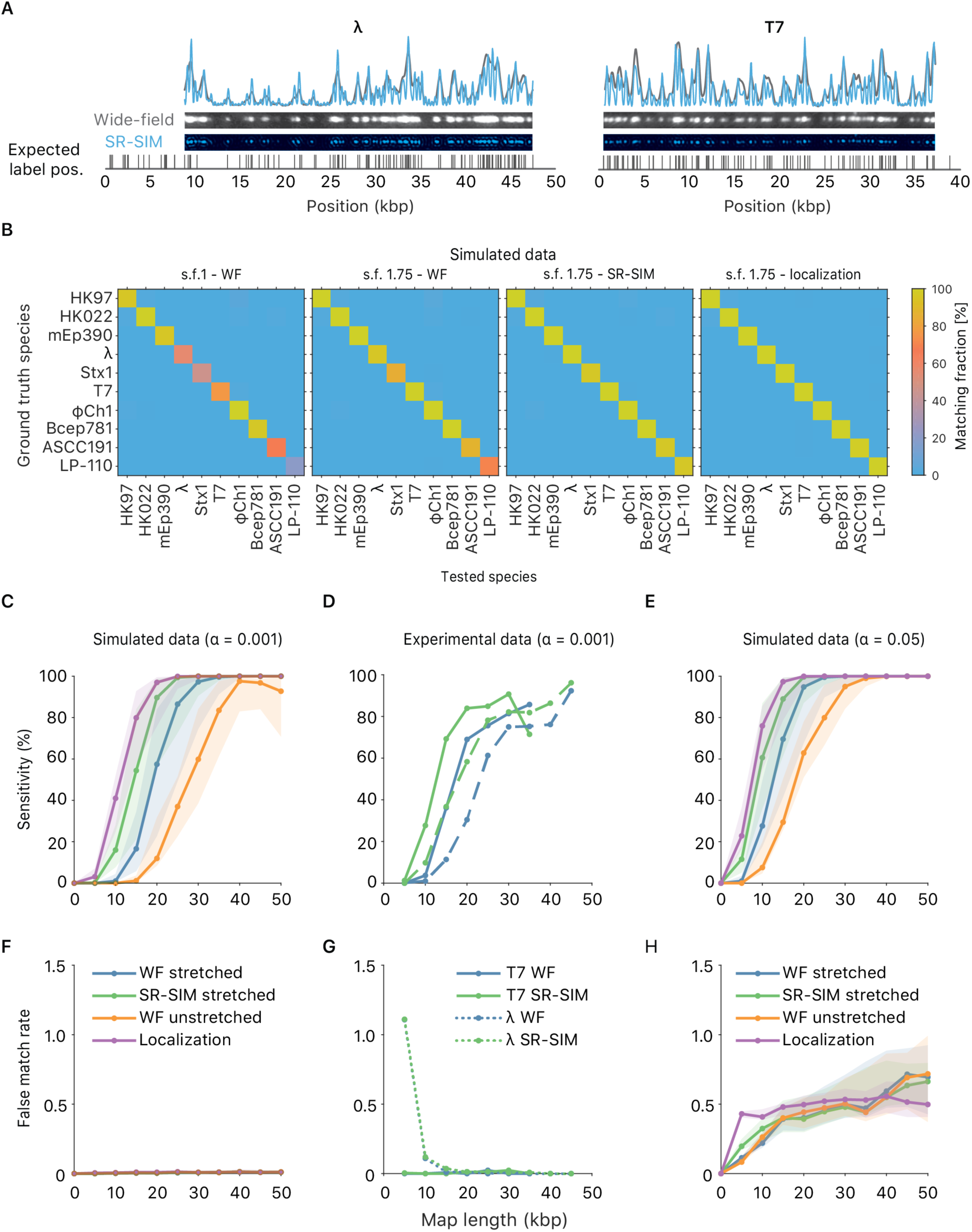
(A) Examples of experimental microscopy images of bacteriophage lambda and T7 DNA fragments, obtained in wide-field (blue) and by SR-SIM (green). The corresponding fluorescence intensity traces are shown at the top. The traces are placed at the location of the genome found by maximizing the cross-correlation. The black vertical lines below the traces indicate the expected dye positions (i.e. the locations of the recognition sequence in the full genome). (B) Assignation matrices for 1000 simulated DNA fragments/species drawn from the full genome of 10 different bacteriophage species and matched to the same 10 species. Significance threshold *α*_1_ = 0.001. Different methods for collecting the DNA fragment measurements are compared. From left to right: Unstretched DNA fragments imaged by wide-field microscopy; Overstretched DNA fragments (stretching factor 1.75) imaged by wide-field microscopy; Overstretched DNA fragments (stretching factor 1.75) imaged by SR-SIM microscopy; Overstretched DNA fragments (stretching factor 1.75) imaged by localization microscopy. See Supplementary Information, Section 3.3, for more detailed versions of the assignation matrices. (C) Simulated data: Bacteriophage identification sensitivity as a function of simulated DNA fragment length. Significance threshold *α*_1_ = 0.001. Solid lines indicate the median over all the 10 species. The shaded areas are circumscribed by the 25-th and the 75-th percentile of the sensitivity values obtained for the set of 10 different species. (D) Experimental data: Sensitivity as a function of DNA fragment length. (E) Simulated data: Sensitivity as a function of simulated DNA fragment length (*α*_1_ = 0.05). (F) Simulated data: False matching rate as a function of simulated DNA fragment length (*α*_1_ = 0.001). The shaded areas are circumscribed by the 25-th and the 75-th percentile of the false matching rate values obtained for the set of 10 different species. (G) Experimental data: False matching rate as a function of DNA fragment length. (H) Simulated data: false matching rate as a function of simulated DNA fragment length (*α*_1_ = 0.05).

The improvement due to overstretching is clearly visible from the number of significant matches on the diagonal (i.e., the true positive rate or sensitivity) of the assignation matrices in Figure 2B (see also Supplementary Information, Section 3.3). In the case of wide-field imaging without DNA overstretching, the sensitivity was rather low, and there was a large variation between species (some species showed around 100% matching sensitivity while some around 20%). Most notably both the species with the lowest labeling density showed low matching sensitivity. DNA overstretching increased sensitivity. Interestingly, imaging unstretched DNA with SR-SIM showed the same improvement of matching sensitivity (not shown) as imaging overstretched DNA with standard wide-field, most likely because the resolution in terms of basepairs is almost identical, which demonstrates that it is such a resolution that determines the sensitivity of the matching. Applying overstretching and increasing the resolution (either by SR-SIM or SMLM) further boosted sensitivity, as shown in the two rightmost matrices in Figure 2B (see also more detailed assignation matrices in Supplementary Information, Section 3.3).

In addition, we found that overstretching and improved resolution reduced the DNA fragment length required to achieve a given level of sensitivity. Hence, shorter DNA fragments can still be correctly identified with high sensitivity. This effect can be seen in Figure 2C where the sensitivity is plotted for different simulated fragment lengths. The improved resolution lowered the fragment length required for maximal sensitivity from 40 kbp down to 20 kbp. Interestingly, while SMLM did improve the matching sensitivity, its improvement was smaller than the improvement of SR-SIM over wide-field, shifting the curve leftward by just a few kbp. Possibly, the sensitivity cannot be improved much by SMLM because not much additional information is revealed by further narrowing the PSF.

Moreover, note that there was a rather large variation in sensitivity across the different species, as can be seen from the shaded areas in Figures 2C and 2E. The shaded areas are circumscribed by the 25-th and the 75-th percentile of the sensitivity values obtained for the set of 10 different species. This variation seems to be mainly due to the variation in labelling density across species. Both species with high labeling density (ΦCh1 and Bcep781) showed the highest sensitivity in matching while species with low labeling density (ASCC191 and LP-110) showed the lowest sensitivity in matching. This dependence of the matching sensitivity on labeling density is most likely due to the ratio between false labels and real labels, as we kept the false-label rate (the number of false labels per kbp) constant in our simulations. Under this condition, DNA optical maps proceeding from genomes with low recognition site density tend to be characterised by a reduced number of fluorescent labels and, therefore, to be less specific than those belonging to species whose recognition site density is higher (for a given map length and a given enzymatic labelling efficiency). It is then reasonable to think that variations like false labels affect the assignation accuracy more for the former than for the latter. And this constitutes a natural drawback intrinsic to the nature of the specific genetic material at hand. The most immediate solution to overcome such a limitation is to guarantee for all the microbial species under study a sufficiently high enzymatic labelling efficiency such that all the analyzed DNA optical maps contain information as specific as possible for a reliable and, possibly, unique assignation. This is evident from Supplementary Figure 13, which suggests that a minimal labeling efficiency of 70% was needed for achieving good matching sensitivity with overstretched DNA images collected by wide-field microscopy. As for fragment length, an increase in resolution also relaxes this requirement to about 60%.

To confirm these effects experimentally we imaged overstretched DNA fragments from bacteriophage T7 and lambda using a SIM microscope, which allowed us to obtain both wide-field and SR-SIM images for each DNA fragment (see Materials and Methods section). The resolution improvement is apparent from the experimental data shown in Figure 2A, where wide-field and SR-SIM data are shown side-by-side. We assessed the resolution improvement experimentally to be close to 2 (see Supplementary Information, Section 2.3). Our experiments showed similar improvements in sensitivity in the experimental data (Figure 2D) as we have seen in the simulated data. Moreover, to investigate the effect of fragment length, the measured DNA maps were cut to various lengths *in silico*. Again, we observed that increasing the resolution lowered the requirement on the DNA size for both lambda and T7.

A higher sensitivity can also be achieved artificially by raising the significance threshold (Figure 2E). However, this improvement comes at the cost of a higher false matching rate (i.e., the number of significant matches to the wrong species – Figure 2H). In contrast, the increased sensitivity obtained by improving resolution does not suffer from this trade-off: the false matching rate is rather independent from resolution. This observation is valid both for strict (*α*_1_ = 0.001 – Figures 2F and 2G) and less strict (*α*_1_ = 0.05 – Figure 2H) significance thresholds. Notice that the significance threshold can anyway be tuned in order to achieve the best compromise between assignation sensitivity and false matching rate by performing DNA optical mapping experiments encompassing known microbial species and by tools like Receiver Operating Characteristic (ROC) curves. (36)

If we consider these results in the context of using DNA mapping as a tool for identification of microbiome species, it is important to realize that besides sensitivity, the throughput of the method is crucial as well. Because a large number of species is involved in the microbiome, it is important to gather enough data to allow their reliable identification. Typical sequencing approaches yield around one million reads. (37) If the same quantity of optical mapping reads is required (i.e., one million DNA maps), then the acquisition time for a single field-of-view on the microscope (containing typically 100 individual DNA fragments) should take at most a few seconds to realistically acquire enough images in one day. Such acquisition speeds are attainable with the SR-SIM and wide-field analysis, but not with SMLM. Although SMLM achieves the highest sensitivity, the image acquisition and analysis are orders of magnitude slower compared to wide-field and SR-SIM.

### Improving specificity by resampling the highest significant matching score

By expanding the set of 10 bacteriophage species to include 2 species closely related to lambda (having 82 and 69% sequence similarity to lambda, respectively) and 2 species closely related to Stx 1 (having 95% and 85% sequence similarity to Stx 1, respectively), we observed that while an improved resolution still increases the overall identification sensitivity, the false positive rate (i.e., 1-specificity) also increases for the closely related species (see Supplementary Figure 12). Figure 3A shows how closely related the added species are. For a detailed overview of the expanded set of 14 species see Supplementary Table 2.

**Figure 3:**
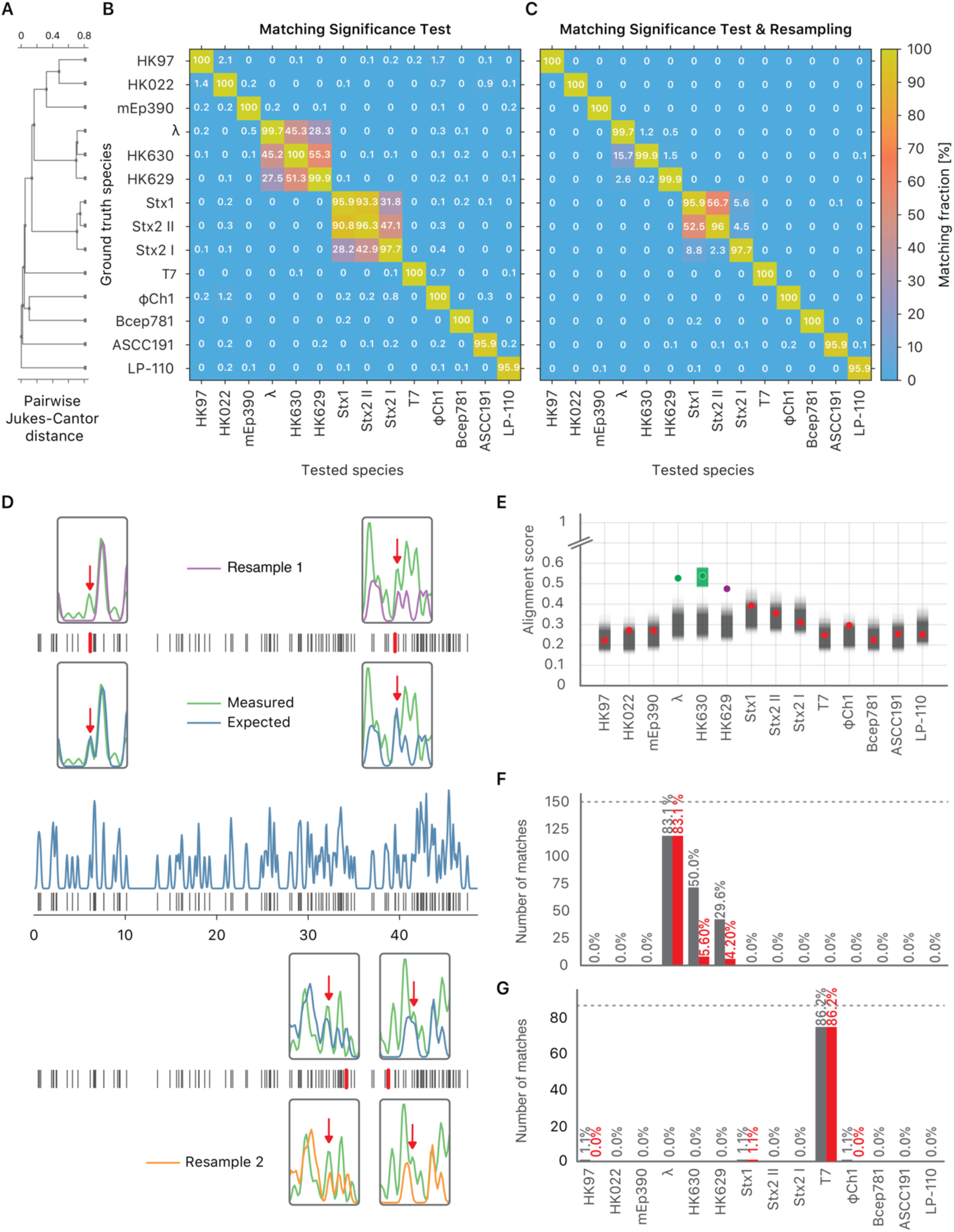
The resampling step improves specificity. (A) Phylogenetic tree of the selected bacteriophages, constructed from the pairwise Jukes-Cantor distance between their sequences. (B) Assignation matrix showing matching percentages yielded by the matching significance test of the matching algorithm for 1000 simulated wide-field data traces per ground truth species. Significance threshold *α*_1_ = 0.001. Note how the regions of confusion between species correspond to short sequence distances in (A). (C) Assignation matrix showing matching percentages yielded by the matching significance test and the resampling step for the same data traces as in (B). Significance threshold *α*_1_ = *α*_2_ = 0.001. Note the reduced confusion in the regions of short sequence distances. (D) Schematic representation of the resampling step. Intensity trace of a measured lambda DNA molecule (green) overlaid with the ideal trace of the same molecule (blue). Underlying dye locations are indicated by black vertical lines. The resampling of the ideal trace is performed by randomly removing two dye locations (red vertical lines) from the matching region (grey box). Two examples of resampled ideal traces are shown (orange and purple). (E) Schematic representation showing the distributions of the maximum cross-correlation scores yielded by the matching significance test and the resampling step, respectively. Experimental data for one measured lambda DNA molecule. The scores for the expected DNA traces are shown by colored dots. The greyscale distributions refer to the randomized scores used for the matching significance test. Red dots indicate non-significant scores (*p*_1_ ≥ *α*_1_). The green and purple dots indicate significant scores (*p*_1_ < *α*_1_). The highest score was found for the tested species HK630 whose ideal trace is therefore resampled within the matching region (green distribution). The score for the tested species lambda was found to be reasonably drawn from the green distribution. The additional match to HK629 can be safely ruled out since its score falls significantly outside the green distribution. The algorithm therefore assigns the DNA map to HK630 and lambda at the same time. (F, grey bars) Results of matching 142 lambda DNA molecules, yielded by the matching significance test (experimental data, SR-SIM microscopy, *α*_1_ = 0.001). The dotted line indicates the total amount of DNA maps concerned. (F, red bars) Results of matching the same molecules, yielded by additionally applying the resampling step (*α*_2_ = 0.001). (G, grey bars) Results of matching 87 T7 DNA molecules, yielded by the matching significance test (experimental data, SR-SIM microscopy, *α*_1_ = 0.001). The dotted line indicates the total amount of DNA maps concerned. (G, red bars) Results of matching the same molecules, yielded by additionally applying the resampling step (*α*_2_ = 0.001).

The assignation matrix for simulated overstretched SR-SIM data in Figure 3B illustrates that, while the sensitivity of the method is high, many false positives are observed within the two sets of closely related species. Therefore, we developed a method to extend the benefits of improved resolution to closely related species as well. Lowering the significance threshold does not solve this issue, as the improved specificity comes at the cost of reduced sensitivity. An alternative approach to improve the specificity of our method, without sacrificing sensitivity, would be to directly compare the matching scores found to be statistically significant by the matching significance test. Ideally, the highest of these matching scores reflects the true positive match. However, because of imperfect enzymatic labeling, the observed score may be lower than in ideal conditions and could vary from case to case. Because of this, closely related species with similar sequences might show very similar matching scores. Therefore, we can think of assigning only to species whose matching score is significantly higher than the others. Ideally, we would like to know how large the spread on the highest matching score due to the labeling uncertainty is. Knowing the spread, we could then test which matching scores are significantly lower than the highest one and reject them.

To incorporate the effect of an imperfect labeling efficiency we emulated the spread on the highest matching score by randomly removing two chosen labels from the corresponding expected DNA map, and re-calculating the matching score (illustrated in Figure 3D). Repeating this procedure many times yielded a distribution for the highest significant matching score. We then tested which of the other species yielded matching scores that were significantly lower and could therefore be rejected (Figure 3E). The method is described in more detail in the Materials and Methods section. In principle, this second computational step might encompass more rigorous computational steps for the estimation of the uncertainty associated to the global cross-correlation function maximum. Nevertheless, defining an accurate null-model accounting for all the different physico-chemical phenomena involved in the generation of a DNA optical map is not straightforward when real-world case-studies are dealt with. Sample heterogeneity and variability are factors which are not easy to control and their effect on the nature and quality of the collected data is considerable and difficult to forecast or infer a priori in complex biological scenarios. In such situations, more elaborated algorithmic methodologies could easily generate overfitting and return unreliable outcomes, especially if their single underlying operations are not exact or are affected by an intrinsic bias resulting from an ill-conditioned theoretical description of the investigated system. This was the rationale behind the way the second level of the presented data analysis technique was conceived: the use of a simpler, more immediate and purely data-driven approach, which may guarantee a sufficient robustness towards the influence of the aforementioned factors. The reported simulated and experimental examples clearly show the great potential of such a proposal in this sense. Furthermore, the selection of the number of fluorescent labels to remove at each resampling iteration is not crucial from a practical point of view. In fact, the second empirical test of the implemented strategy simply provides a refinement of the assignation results yielded by the first one and is nested to them. That is to say that, if the estimation of the uncertainty on the global cross-correlation function maximum does not lead to a reduced assignation ambiguity, the output proceeding from the first step of the statistical workflow (which is anyway meaningful from the identification point of view) will be retained. All this renders a remarkably good compromise between computational complexity and efficiency and microbial species differentiation accuracy.

Figure 3E illustrates this procedure on experimental data taken from a lambda DNA molecule imaged by SR-SIM. In this example, significant matching scores are found for three closely related species: lambda, HK 630 and HK629. The highest matching score is found for the (wrong) species HK630. Resampling the highest matching score and testing which scores are significantly lower allows rejecting the match to HK629, but not lambda. Hence, the resampling procedure has improved specificity by rejecting one false positive, without rejecting the true positive (lambda).

Our method for the identification of imaged DNA molecules now consists of two steps. First, a matching significance test is performed on the matching score and all non-significant matches are rejected. Second, in the resampling step, all significant matches with matching scores significantly lower than the highest significant one are rejected. As can be seen from the results for the simulated overstretched SR-SIM data in Figure 3C, applying the resampling step clearly improved specificity, while having only a minor effect on sensitivity. We observed the same improvement in experimental data from bacteriophages lambda (Figure 3F) and T7 (Figure 3G).

One might wonder whether it is necessary to perform the initial matching significance test at all if the resampling step allows distinguishing between species on its own. Indeed, if the reference database of tested species contains all possible species present in the sample, this approach would yield the same results. However, if no matching score significance test is carried out, the resampling step will always return at least the species with the highest matching score. Thus, if the sample under study contains species that are not present in the reference database, measured maps from such unknown species will always be assigned to wrong species. The matching score significance test is therefore required to eliminate false matches for unknown species. Considering the lack of assembled genomes for a lot of species in the microbiome (colloquially known as the dark matter of the microbiome) it is necessary to perform a significance test before comparing matching scores.

### Simulated identification of bacterium-sized genomes

To investigate if the identification performance varies when the reference species have a much longer genome (a bacterial genome, for example, is about 2 orders of magnitude larger than the one of a bacteriophage), we created artificially large genomes *in silico* by inserting the sequence of a bacteriophage (lambda and T7) in the middle of the sequence of a bacterial genome (*Bacteroides thetaiotaomicron* VPI-5482) having similar labeling density (lambda: 2.5/kb, T7: 2,8/kb, bacteroides: 2.8/kb). Next, we used our experimental SR-SIM data for lambda and T7 DNA fragments and matched them to the artificial bacteria, as well as to the bacterium without the inserted phage DNA. We also matched to two other bacteria as a control: *Lactobacillus Reuteri* and *Escherichia Coli* (strain K-12 substrain MG1655). After performing the matching significance test, we found that matching sensitivity to the artificial bacterium was lower than to the phage itself (both for phages lambda and T7, as shown in Figures 4A and 4B). We were able to bring the sensitivity for the artificial bacteria back towards the level of phages by implementing a local normalization on the expected maps before the calculation of the cross-correlation function. The rationale behind the local normalization was that local regions of high labelling density bias the cross-correlation to high values, causing false matches. However, these false matches did not pass the matching significance test, since the reshuffled bacterial genomes also contained random regions of high local labelling density, which yielded high randomized matching scores. By locally centering and standardizing the expected DNA maps in a 5kb window, high cross-correlation values in high local density regions were avoided. With local normalization, matching to the artificial bacteria performed similarly as when matching to the phages themselves, as shown in Figures 4C and 4D.

**Figure 4:**
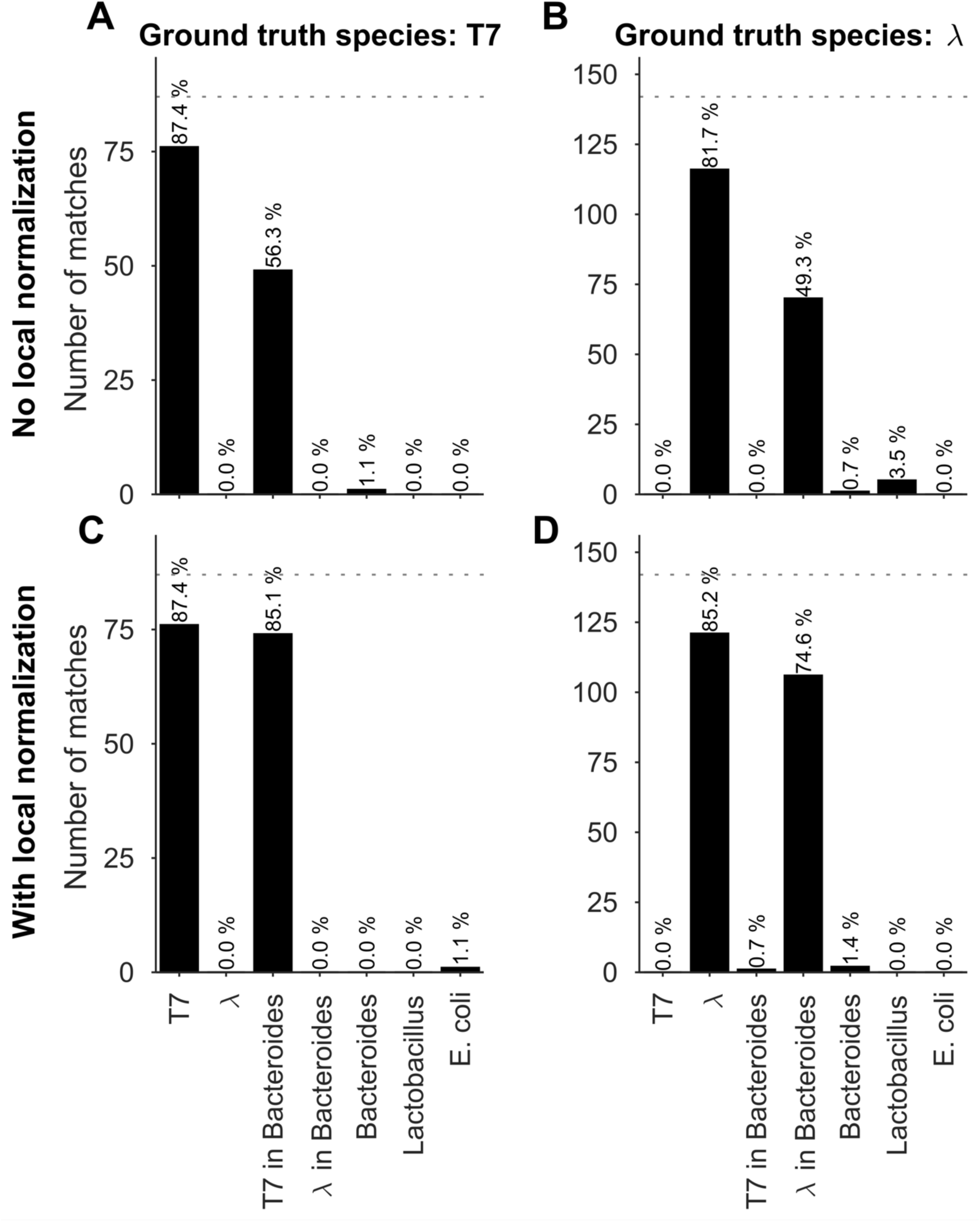
Identification of bacteria simulated by using experimental bacteriophage data recorded with SR-SIM. Results of matching bacteriophage T7 and lambda DNA molecules to phage and artificial bacterial genomes, yielded by matching significance testing (*α*_1_ = 0.001). (A) Ground truth species: T7, no local normalization. (B) Ground truth species: lambda, no local normalization. (C) Ground truth species: T7, 5kb-window local normalization. (D) Ground truth species: lambda, 5kb-window local normalization.

### The case of *V. Harveyi*

In order to prove the usefulness and suitability of the proposed methodology for the assessment of more complex biological samples, 601 optical maps of *V. Harveyi* DNA were recorded based on the experimental procedure described in the Materials and Methods section and fed to the developed algorithm.

*V. Harveyi* can be regarded as a naturally “calibrated” biological system. Every *V. Harveyi* cell, in fact, contains DNA constituted by 2 single chromosomes (chromosome #1: accession number CP000789.1, size 3.77 Mb – chromosome 2: accession number CP000790.1, size 2.20 Mb) and a varying amount of plasmids (accession number CP000791.1, size 0.09). As the ratio of occurrence of these 2 chromosomes does not change across cells, its expected value can be easily estimated as the ratio of their size, which is equal to 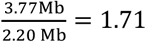. Assuming that the imaged optical maps cover more or less homogeneously this full genome and considering the fact that the M.TaqI labeling density inside both chromosomes is approximately the same (4.489 sites/kb for chromosome #1 and 4.654 sites/kb for chromosome #2), the ratio between their relative abundance yielded by the two consecutive algorithmic steps of our assignation strategy should also match 1.71. Figure 5 confirms this point: the calculated ratio between the relative abundance of chromosome #1 (0.59) and chromosome #2 (0.35) equals 1.69, which is closely in agreement with the theoretical expectation. This example shows that the identification procedure described in this article represents a powerful data analysis tool which might aid the resolution of mixtures of sequenced species characterized by single DNA molecule fluorescence optical mapping. Given the higher level of noise observed for this particular case-study, *α*_1_ and *α*_2_ were here set to 0.05 and 0.001, respectively. A 5kb-window local normalization was also carried out.

**Figure 5:**
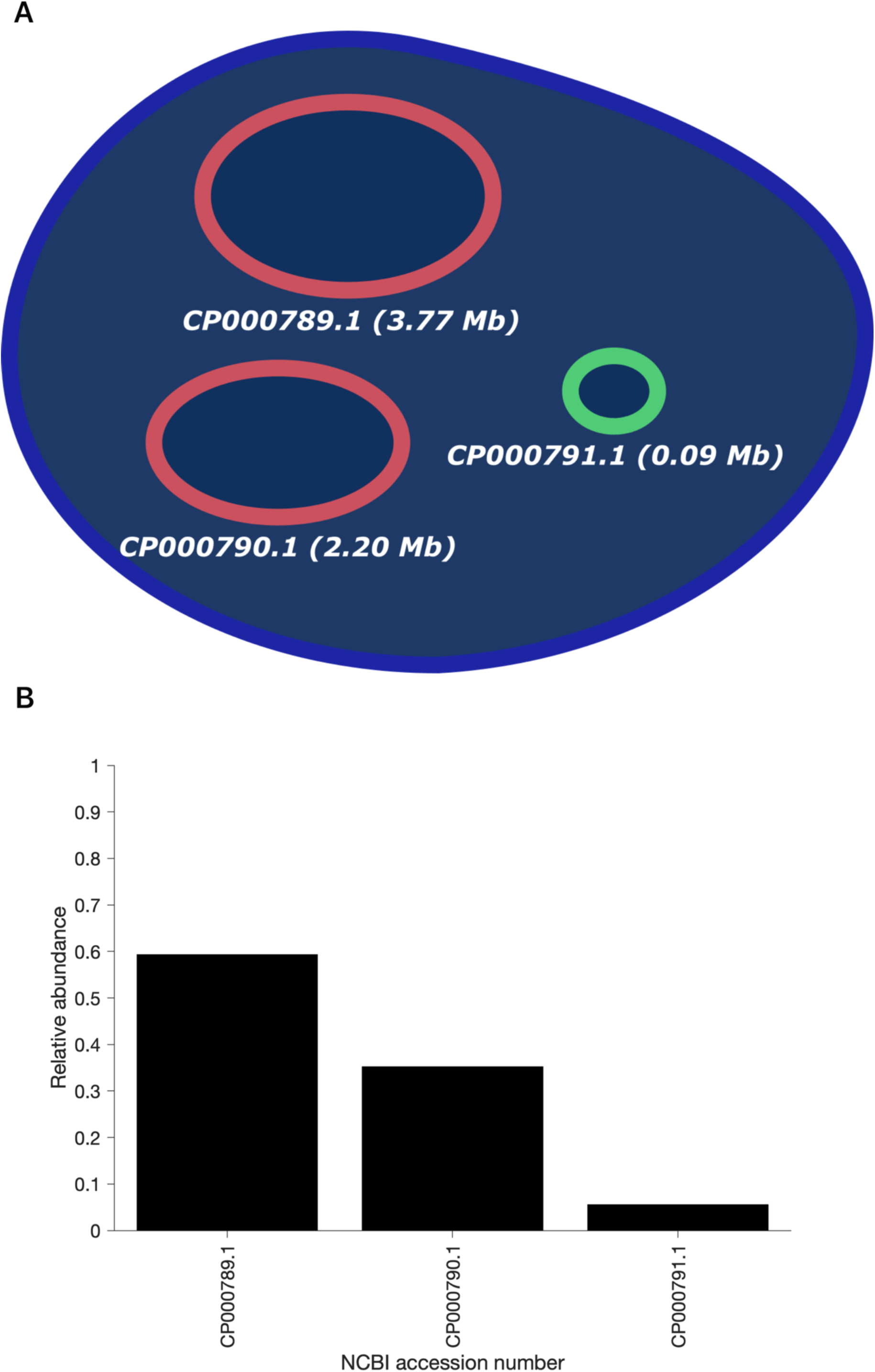
(A) Schematic representation of the genetic content of *V. Harveyi*. (B) Abundance of chromosome #1 (accession number CP000789.1), chromosome #2 (accession number CP000790.1) and plasmids (accession number CP000791.1) relative to the total amount of assigned optical maps sampled from *V. Harveyi* DNA (*α*_1_ = 0.05, *α*_2_ = 0.001, 5kb-window local normalization). The results reflect the expected occurrence of the three different constituents.

## CONCLUSIONS

In this article, we proposed an improved method for identifying microbial species based on single-molecule double-stranded DNA maps. We based ourselves on intensity profiles extracted from microscopy images. These intensity profiles can be compared to a reference sequence by generating an expected map and cross-correlating it with the measured map. More specifically, we first subject the retrieved alignment to a matching significance test, where we calculate an empirical *p*-value for the alignment. We showed, for this matching significance test, how an increase in resolution, both through super-resolution microscopy as well as overstretching, improved identification sensitivity. This means many of the false assignations are filtered out. However, it does not remove assignation ambiguity for closely related species. For this reason, we have developed a second algorithmic step, where we resample the highest significant matching score by generating variations of the expected map with missing labels (i.e., mimicking lower labeling efficiency). This resampling step assigns the map to a single species if its matching score is significantly higher than the significant matching scores for other species. Importantly, the method we proposed for species identification is independent from the tool used to extract the DNA maps and could easily be applied to other datatypes such as current traces through a nanopore. Finally, we showed that, with the addition of a local normalization procedure, this method can also be extended to identification of bacterial species. As an alternative for resampling the theoretical traces, one could also generate surrogates of the acquired intensity profiles using the Fourier phase randomization method that preserves the correlation structure of the empirical data. (38) Recent extensions of this method to graph structures (39) could also be considered if additional information about local stretching are available.

It is fundamental to notice that testing the statistical significance of the matching score allows the whole proposed approach to be robust against biased definitions of the database of reference species. This is a clear advantage over just assigning these fragments to the species for which a maximum matching score is found as it reduces the impact of false positives on the final outcomes. Since currently not all microbial species are sequenced, unknowns are to be expected in real world scenarios. Therefore, not assigning a DNA map will be important when matching the experimental maps against a sub-database containing only the species of interest (since matching against all known species would be computationally very demanding). Such scenarios will occur frequently when studying the change of composition of a few species in a highly heterogeneous system such as the gut microbiome. For all these reasons and considering the possibility our proposal offers of easily recognizing common genomic subsequences shared by multiple species, we believe this novel methodology can be particularly suitable for handling even very complex real case-studies.

Finally, when creating an abundance profile of the microbiome, high sample throughput is critical for acquiring enough data. Localization microscopy is known for requiring a lot of time for a single image, whereas SR-SIM is a lot faster, making SR-SIM a more realistic imaging tool for obtaining enough DNA maps. A second important element is the analysis of the DNA maps. Analysis of shotgun metagenomic reads from microbiome samples is typically very challenging from a computational point of view due to the large quantities of data involved. (37) This issue also applies to optical mapping and the method presented here: all measured DNA maps need to be aligned to the expected DNA maps for all target species. Nevertheless, the selection of the reference DNA sequences is actually not a critical step. The algorithmic procedure proposed here, in fact, performs a separate test for every single species under study in the attempt of assessing whether a particular DNA map belongs to its corresponding genome or not. This way, one can easily reduce the database constructed for identification purposes so as to encompass only few microorganisms of interest while guaranteeing high robustness against optical maps proceeding from unknown species (that, as specified before, ideally would not be assigned). This would dramatically decrease the computational load and cost in real-world scenarios characterised by the presence of unsequenced or partially sequenced genomes and extreme complexity and heterogeneity (e.g., gut microbiome mapping) without jeopardizing the identification quality and minimizing the number of false assignations. Moreover, alignment of DNA maps by cross-correlation can be implemented very efficiently by converting the signals into the Fourier domain. Due to the convolution theorem, the cross-correlation then becomes a computationally cheaper multiplication. Moreover, the database of expected DNA maps for all target species needs to be Fourier-transformed only once, after which it can be re-used for each alignment. As an example, a single cross-correlation alignment against a 6.3 Mbp bacterium (*Bacteroides thetaiotaomicron* VPI-5482) took 86 ms, whereas a dynamic programming algorithm took 3.3 seconds (see Supplementary Figure 17 for more details), almost a 40-fold increase. Additionally, cross-correlation analysis generated more correct significant matches compared to dynamic programming alignment (see Supplementary Figure 16).

## ACKNOWLEDGEMENTS

The authors would like to thank Marcel Müller, Wim Vandenberg and Linda Mhalla for helpful discussions. The Zeiss SIM Elyra microscope was acquired through a CLME grant from Minister Lieten to the VIB BioImaging Core. The Tesla K40 GPU used for this research was donated by the NVIDIA Corporation.

## Footnotes

i If one assumes labelling to be a Poisson process, the probability of finding one or more extra fluorescent labels within a PSF is given by: 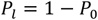 where 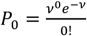 based on the definition of the Poisson distribution. *v* here denotes the average expected labelling rate within a PSF, yielded by the size of the PSF divided by the expected distance between two labels on a random string (4^4^ = 256 bp). Expressing the FWHM of the PSF in basepairs as: 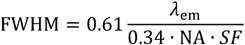 with *λ*_em_ = 576 nm and NA = 1.4 as described before, *SF* being the DNA stretching factor and 0.34 representing the DNA basepair distance, it subsequently holds that for unstretched DNA (*SF*= 1): 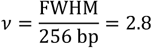 and 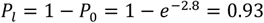 *quod erat demonstrandum*.

