## Supporting information for "Identifying microbial species by single-molecule DNA optical mapping and resampling statistics"

Supplementary Information

### 1. Bacteriophage species

| Bacteriophage name | Shorthand name used in figures | NCBI accession number | Sequence length (bp) | Number of recognition sites | Density of sites (sites/kb) |
|---|---|---|---|---|---|
| **Escherichia virus Lambda** | λ | J02459 | 48502 | 121 | 2,49 |
| **Escherichia virus HK97** | HK97 | NC_002167 | 39732 | 147 | 3,70 |
| **Escherichia virus HK022** | HK022 | NC_002166 | 40751 | 165 | 4,05 |
| **Enterobacterial phage mEp390** | mEp390 | NC_019721 | 40029 | 154 | 3,85 |
| **Escherichia Stx1 converting phage** | Stx1 | NC_004913 | 59866 | 139 | 2,32 |
| **Enterobacteria phage T7** | T7 | V01146 | 39937 | 111 | 2,78 |
| **Lactococcus virus ASCC191** | ASCC191 | NC_017688 | 32414 | 33 | 1,02 |
| **Listeria phage LP-110** | LP-110 | NC_021785 | 65132 | 74 | 1,14 |
| **Natrialba phage PhiCh1** | Φ Ch1 | NC_004084 | 58498 | 1048 | 17,92 |
| **Burkholderia virus Bcep781** | Bcep781 | NC_004333 | 48247 | 528 | 10,94 |

**Table 1:** Set of 10 bacteriophages used to generate simulated data and taken as target species. The set of species includes bacteriophages lambda and T7 (orange text), for which we also obtained experimental data. The rows with blue text refer to the two species that were added to investigate the effect of a low density of recognition sites. The rows with red text refer to the two species that were added to investigate the effect of a high density of recognition sites. Bacteriophage lambda and the species indicated in black characters are from the Siphoviridae family.

| Bacteriophage name | Shorthand name used in figures | NCBI accession number | Sequence length (bp) | Number of recognition sites | Density of sites (sites/kb) |
|---|---|---|---|---|---|
| **Enterobacteria phage HK630** | HK630 | NC_019723 | 47090 | 122 | 2,59 |
| **Enterobacteria phage HK629** | HK629 | NC_019711 | 47288 | 120 | 2,54 |
| **Escherichia phage Stx2 II** | Stx2 II | AP005154 | 62706 | 157 | 2,50 |
| **Stx2 converting phage I** | Stx2 I | AP004402 | 61765 | 151 | 2,44 |

**Table 2:** Set of 4 additional bacteriophages used to generate simulated data and taken as target species. The first two species (green text) were chosen because they are closely related to bacteriophage lambda, the second two (purple text) because they are closely related to Stx 1.

### 2. Experimental SR-SIM

#### 2.1 Scanning the stretching factor

During the calculation of the matching scores, the overstretching factor was allowed to vary from 1.7 to 1.76 in steps of 0.01. A broader and coarser scan of the overstretching factor was also performed. This revealed the optimal overstretching factors to be indeed in the 1.7 – 1.76 range. The histograms in Figure 1 show the optimal stretching factor for DNA fragments measured by the SIM microscope. Only significant matches ($\alpha_1 = 0.001$) are shown. The optimal stretching factor was found by maximizing the matching score as described in the Materials and Methods section of the main text of this publication (see Section "Calculation of the matching score").

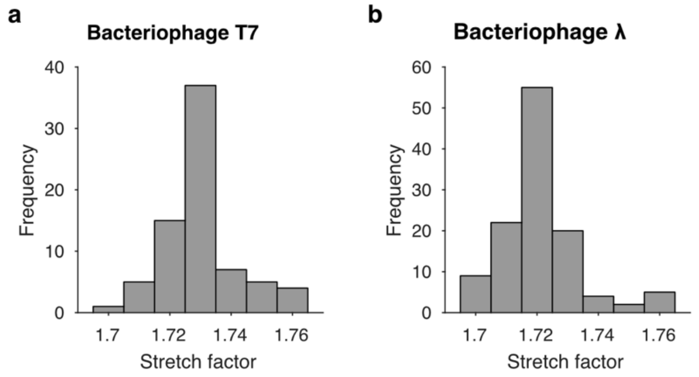


**Figure 1:** Optimal overstretch factors for DNA fragments included in the experimental SR-SIM dataset (significant matches only, $\alpha_1 = 0.001$). (a) Bacteriophage T7, 82 significant matches (out of 87 DNA fragments). (b) Bacteriophage lambda, 132 significant matches (out of 142 DNA fragments).

### 2.2 Length histograms of experimental data

The length histograms in Figure 2 show the DNA fragment lengths for the experimental data measured by the SIM microscope. Each DNA fragment length was first measured in camera pixels by manually segmenting the images by ImageJ and was then converted to basepairs using the optimal overstretching factor estimated as described before.

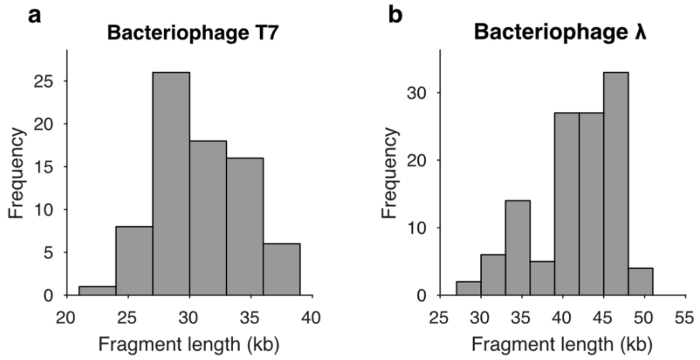


**Figure 2:** Length histograms for DNA fragments included in the experimental SR-SIM dataset (significant matches only, $\alpha_1 = 0.001$). (a) Bacteriophage T7, 82 significant matches (out of 87 DNA fragments). Full-length T7 DNA: 39.9 kb. (b) Bacteriophage lambda, 132 significant matches (out of 142 DNA fragments). Full-length lambda DNA: 48.5 kb.

### 2.3 Determining the optical resolution

The DNA fragments are much narrower than the width of the microscope Point Spread Function (PSF). Hence, the PSF Full-Width at Half-Maximum (FWHM) can be measured perpendicularly to the length of the DNA fragment. We first averaged each DNA fragment along its length to get an average intensity profile perpendicular to it. This intensity profile was fitted with a Gaussian function. The expected FWHM for wide-field is

$$\mathrm{FWHM} = 0.61\frac{\lambda_{\mathrm{em}}}{\mathrm{NA}} = 251\ \mathrm{nm},$$

where $\lambda_{\mathrm{em}}$ is the emission maximum wavelength of the fluorescent label (Rhodamine B, $\lambda_{\mathrm{em}} = 576$ nm), and NA is the microscope objective numerical aperture (NA = 1.4). For SR-SIM, the expected FWHM is about half the wide-field value. The histograms in Figure 3 show the measured FWHMs on the SIM microscope in wide-field and SR-SIM modes, for bacteriophages lambda and T7. It can be seen that the expected FWHMs correspond quite well to the measured values for both wide-field and SR-SIM.

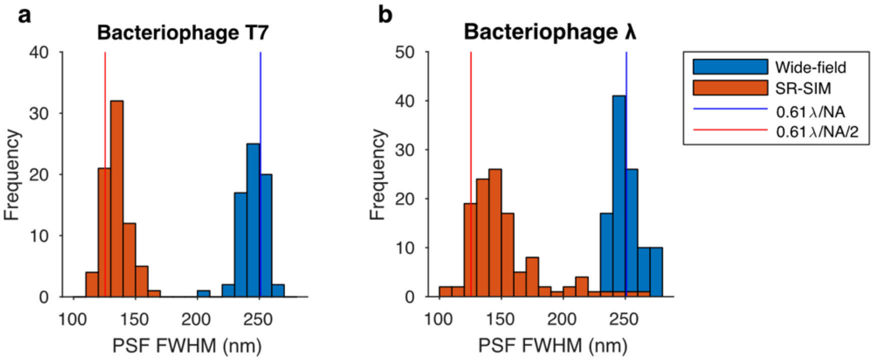


**Figure 3:** FWHM of the PSF measured perpendicularly to the DNA fragments included in the experimental wide-field and SR-SIM datasets (significant matches only, $\alpha_1 = 0.001$).

### 2.4 Pseudo-Receiver Operating Characteristic (pseudo-ROC) curve analysis

Receiver Operating Characteristic (ROC) curves can be used to investigate the identification performance of the matching significance test of our method, independently of the user-defined significance threshold. ROC curves display the true-positive identification rate (TP rate = sensitivity) against the false-positive identification rate (FP rate = 1-specificity) at different significance threshold settings.

Nevertheless, it should be noticed that our method does not allow arbitrarily low significance thresholds to be used, as the smallest $p_1$-value that can be calculated is

$$p_{1,\min} = \frac{1}{N_p + 1}$$

where $N_p$ is the number of label position permutations. Hence, if a lower significance threshold is needed, more permutations should be carried out. The lowest meaningful $p_1$-value threshold that can be considered is therefore limited by the amount of processing time available.

Figure 4 shows the improved identification performance for SR-SIM compared to wide-field data (the former lead to a larger area under the pseudo-ROC curve which constitutes a measure of the overall identification accuracy). However, for closely related species (Figure 4d, inset), increasing resolution does not provide any clear improvement

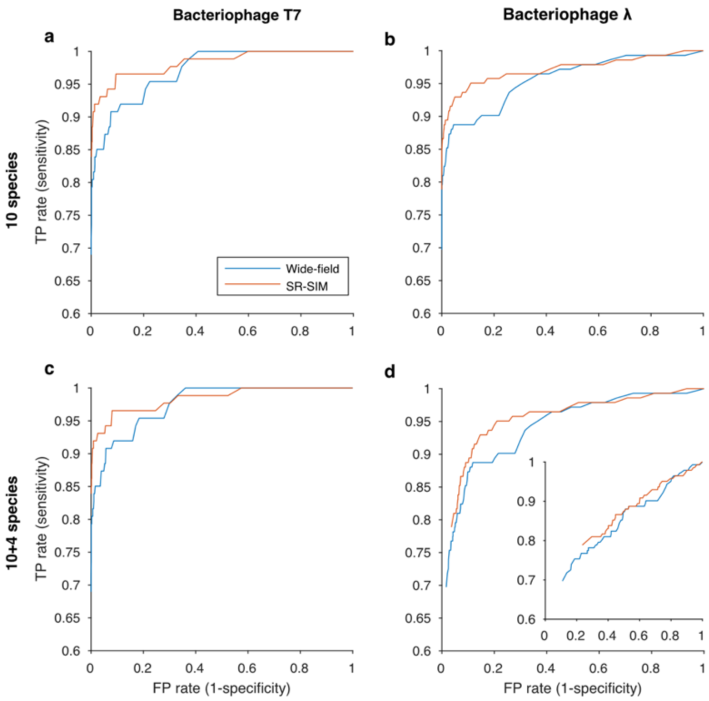


**Figure 4:** pseudo-ROC curves for experimental wide-field and SR-SIM data of bacteriophage T7 (a, c) and bacteriophage

lambda (b, d), obtained using only the matching significance test of the identification method. For (a, b), the set of tested species consists of the 10 species in Table 1. For (c, d), the set of tested species consists of the 10 species in Table 1 plus the 4 species in Table 2. The inset in (d) shows the pseudo-ROC curve obtained when the set of tested species consists only of bacteriophage lambda, and its 2 closely related species: Enterobacteria phage HK630 and Enterobacteria phage HK629.

Note that the resampling step of our bacteriophage identification method introduces a second $p$-value ($p_2$) and improves specificity, but not sensitivity. As the outcomes resulting from it are nested to those proceeding from the matching significance test, ROC curves are not the best way to investigate its performance.

#### 2.5 Dry versus Milli-Q samples

We compared samples of overstretched DNA on coverslips with Milli-Q water on top to dry samples. The top-side of the samples was carrying the labeled DNA. The oil-immersion microscope objective was below the coverslip (inverted microscope).

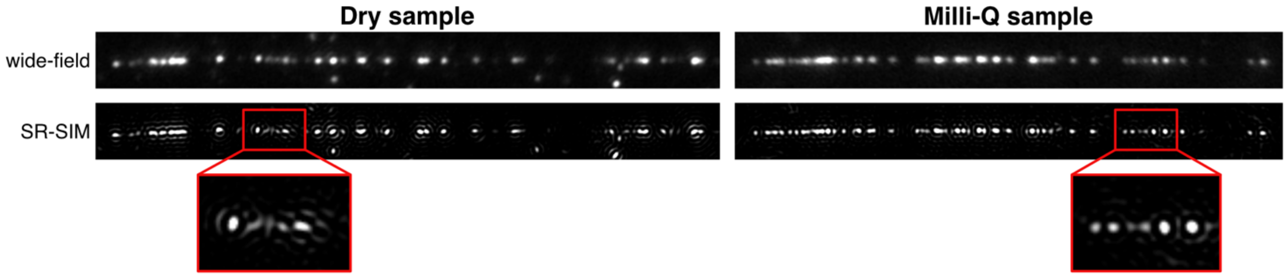


**Figure 5:** Comparison of wide-field and SR-SIM images of lambda DNA in dry and Milli-Q samples.

Figure 5 shows a comparison of typical DNA fragments imaged in wide-field and SR-SIM modes for dry and Milli-Q samples (not the same DNA fragment). Note the elongated shapes of the PSFs in the SR-SIM image of the dry sample (Figure 5, bottom left). These elongated shapes were much less apparent in the SR-SIM image of the Milli-Q sample (Figure 5, bottom right). In the wide-field images, no differences were readily observable.

In order to investigate the effect of the differences between dry and Milli-Q samples on the bacteriophage identification performance, we applied the matching significance test and the resampling step to both types of samples. We noticed an improved sensitivity for the Milli-Q samples (see Figure 6). This improvement was observed for both the wide-field and the SR-SIM data.

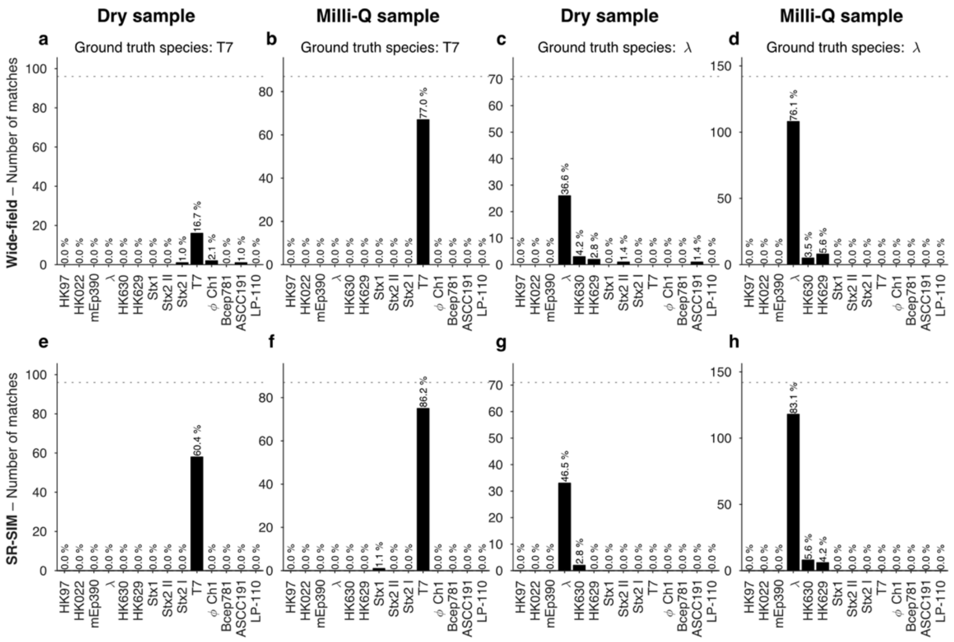


**Figure 6:** Comparison of the identification performance (matching significance test and resampling step) for dry samples

and Milli-Q samples. (a)-(d) wide-field data. (e)-(h) SR-SIM data. $\alpha_1 = \alpha_2 = 0.001$. The dotted line indicates the total amount of DNA maps concerned. Sensitivity was found to be much higher for Milli-Q samples. The set of tested species consisted of the 10 species in Table 1 plus the 4 species in Table 2.

We hypothesize that the improvement in the identification sensitivity might stem from a more uniform fluorescence intensity across the labels. If the 3D orientation of the excitation and emission dipoles of the fluorescent labels is fixed during the camera exposure time, not all labels would be equally bright. However, the matching scores are calculated by cross-correlation with the expected intensity profiles, which are generated by assuming all labels are equally bright. This scenario of uneven dye brightness might apply for the dry samples, while the dipoles in the milli-Q samples might have more rotational freedom, thereby averaging out their brightness during the camera exposure time. A more uniform brightness across labels yields better matching scores and, thus, an improved sensitivity.

Note that the wide-field data was obtained by averaging the 25 frames of the illumination patterns with 5 orientations and 5 phases/orientation of the SR-SIM dataset. The fluorescence excitation light was linearly polarized, perpendicular to the plane of incidence to maximize modulation contrast in the illumination patterns. Hence, averaging over the illumination pattern orientations amounts to averaging 5 angles of linear polarization, each parallel to the sample plane.

In our measurements, the fluorescence excitation polarization was always parallel to the sample plane. Because the fluorescent dipoles might also have an out-of-plane component, a further improvement in dye brightness uniformity might be achieved by adding an out-of-plane component to the excitation polarization.

#### 2.6 Matching positions

Maximizing the cross-correlation also yielded the estimated position of the measured DNA fragments within the genome of the reference species. Figure 7a shows the positions of the bacteriophage lambda DNA fragments within the lambda genome (full length: 48.5 kb) for the various significant matches ($\alpha_1 = 0.001$). As described in the main text, we also matched the experimental DNA fragments to the sequence of an artificial bacterium. The artificial bacterium sequence was composed of the DNA sequence of Bacteroides thetaiotaomicron VPI-5482 with bacteriophage lambda DNA inserted in the middle. The reduction in sensitivity described in the main text can also be observed by comparing Figure 7a and Figure 7b. Because less DNA fragments matched significantly, Figure 7b contains less DNA fragments. The same applies to T7 (Figure 7e and Figure 7f). No significant matches were found outside of the inserted bacteriophage: that is to say the bacteriophage fragments never matched significantly to the bacterium sequence. When applying a local normalization on the expected DNA map of the artificial bacterium, sensitivity was improved (Figure 7c, d and Figure 7g, h). Note that the matching positions are not influenced by the local normalization. The local normalization was performed by subtracting the mean of the expected DNA map within a sliding window, and dividing by the standard deviation of the expected DNA map within the same window. The length of the sliding window was chosen to be 5 kb, as a compromise between not removing too much structure from the DNA maps and avoiding the cross-correlation to be positively biased in regions of high local labelling density.

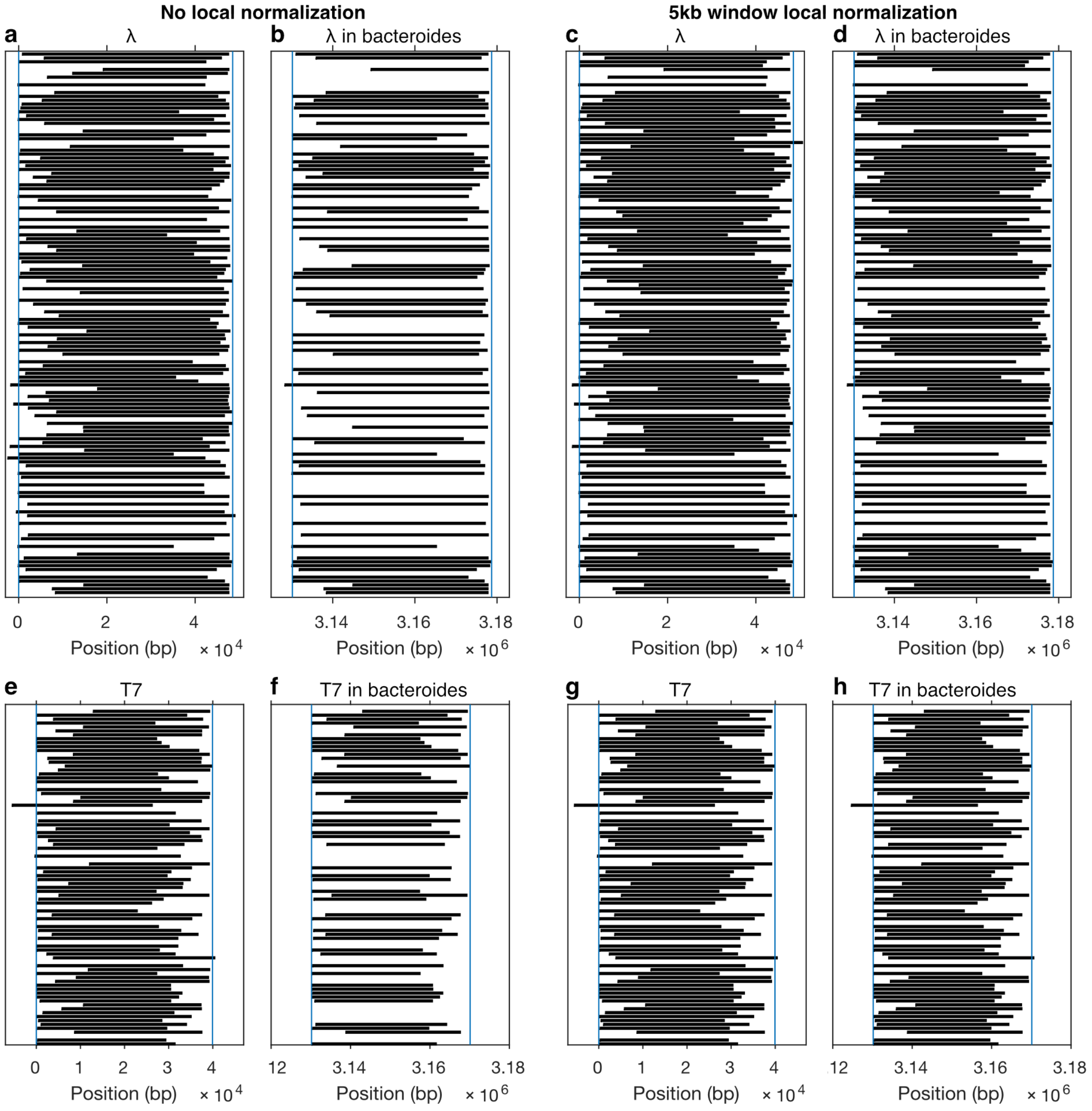


**Figure 7:** Matching positions for experimental SR-SIM data of bacteriophage lambda and T7 DNA. Vertical blue lines indicate the extent of the bacteriophage DNA sequence. Black rectangles represent the length and the position within the target species' genome of significantly matched DNA fragments ($\alpha_1 = 0.001$). (a) Lambda DNA fragments matched to bacteriophage lambda. Matching scores were calculated without local normalization. (b) Lambda DNA fragments matched to Bacteroides thetaiotaomicron VPI-5482 with the DNA sequence of bacteriophage lambda inserted in the middle of its genome (denoted by "lambda in bacteroides"). No local normalization. Only the middle part of the position axis is shown as no significant matches were found to be outside this region. (c) Lambda DNA fragments matched to bacteriophage lambda with a 5kb local normalization window. (d) Lambda DNA fragments matched to lambda in bacteroides with a 5kb local normalization window. (e) T7 DNA fragments matched to bacteriophage T7 without local normalization. (f) T7 DNA fragments matched to T7 in bacteroides without local normalization. (g) T7 DNA fragments matched to bacteriophage T7 with 5kb local normalization window. (h) T7 DNA fragments matched to T7 in bacteroides with 5kb local normalization window.

### 2.7 Fluorescence blinking of dyes on DNA fragments when adding blinking buffer

In order to investigate if blinking-based super-resolution imaging techniques (e.g. Stochastic Optical Fluctuation Imaging, SOFI; or STochastic Optical Reconstruction Microscopy, STORM) could easily be applied in such a scenario, we imaged labeled DNA fragments (bacteriophage T7) in a blinking buffer solution (10 w/v% D(+)-Glucose – Sigma – 0.5 mg/ml Glucose oxidase – Sigma – 34 µg/ml Catalase – Sigma – 142 mM Beta-mercaptoethanol – Sigma – in the presence of 10 mM NaCl – Thermo – and buffered at pH 8 with tris-HCl – Carl Roth). A movie of 900 frames was recorded (50 ms of exposure time per frame). Figure 8 shows the results obtained for a typical DNA fragment. The time-averaged signal for this fragment matched significantly to T7 ($\alpha_1 = 0.001$), but the data were not suitable for blinking-based super-resolution analysis, as the waterfall plot at the lower half of Figure 8 shows. Indeed, some dyes seemed to exhibit blinking, while others did not. Moreover, some dyes seemed to switch between slow and fast blinking. Further research would be needed to investigate this phenomenon. Instead, we chose to use SR-SIM for super-resolved imaging, as this easy-to-use technique does not require blinking of the dyes.

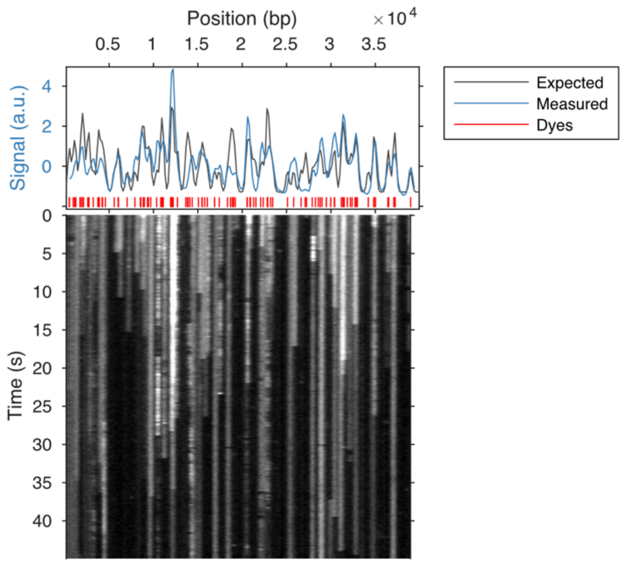


**Figure 8:** A typical bacteriophage T7 DNA fragment imaged in a blinking buffer. Top panel: time-averaged signal (blue line) matched significantly ($\alpha_1 = 0.001$) to the expected signal for T7 (black line). The underlying dye locations are indicated by vertical red lines. Bottom panel: waterfall plot of the fluorescence intensity over time. Note the many different blinking behaviors for the different dyes.

### 3. Simulating optical DNA mapping

#### 3.1 Detailed overview of the simulation model

The different steps of the simulation model we developed for this work are outlined in Figure 9, and listed below.

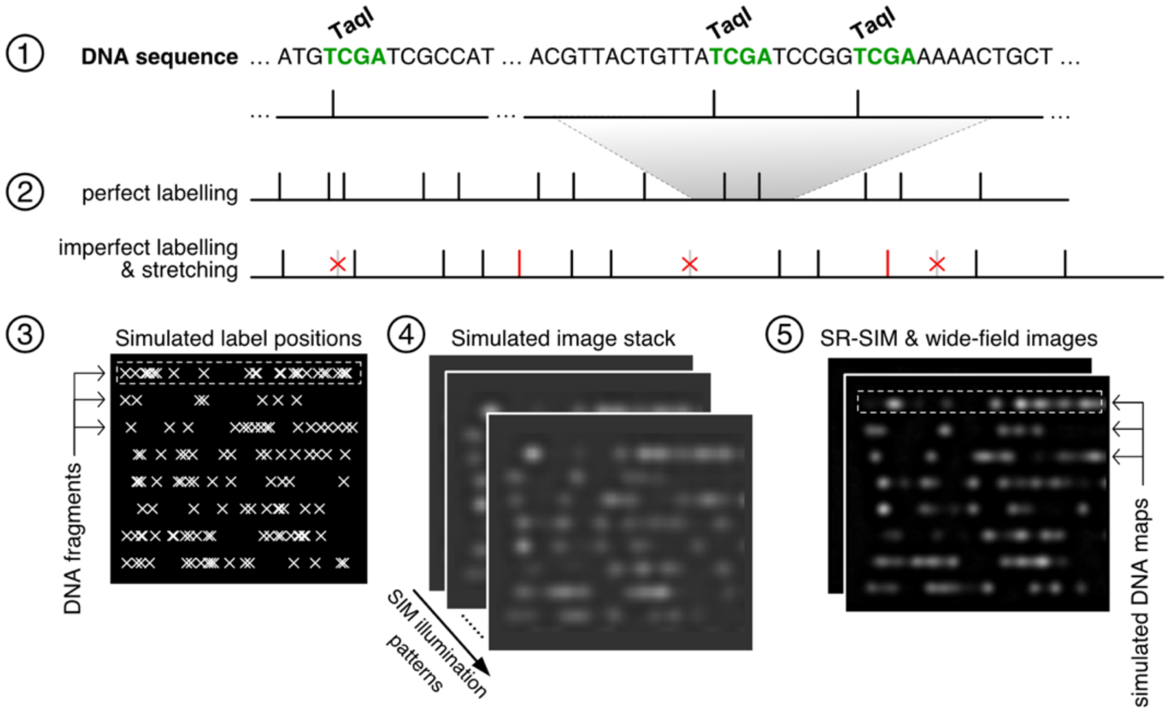


**Figure 9:** Schematic representation of the computational steps of the simulation model.

(1) The DNA sequence of the bacteriophage of interest was downloaded from the NCBI database;

(2) The locations of the recognition sites were listed. In Figure 9, the M.TaqI enzyme recognition site is shown (TCGA). Imperfect labeling was simulated: 70 % labeling efficiency, false positive labeling rate of 0.5 labels/kbp. DNA overstretching was simulated with a factor of 1.75;

(3) 1000 DNA fragments with random starting positions in the bacteriophage DNA, and of length of 35 kb were simulated and placed on horizontal lines of the simulated sample;

(4) The 25 SIM illumination patterns were applied having the same 5 orientations and 5 phases/orientation as in the experiments. The simulated illumination patterns were sinusoidal patterns with a modulation depth of 0.7 corresponding to the value estimated during the reconstruction of the experimental SR-SIM images by the fairSIM algorithm. Random fluorophore bleaching (average bleaching time = 100 frames) and photon noise were simulated. The average number of collected photons per fluorescent label and per camera exposure time (300 ms) was estimated to be $\sim 10^4$. The simulated PSF was a 2D Gaussian with FWHM of $0.61\lambda/\text{NA}$. The fluorescence emission wavelength, $\lambda_{\text{em}}$, was 576 nm, corresponding to the emission maximum of the Rhodamine-B dye used in the experiments. The simulated microscope also had the same configuration as the experimental microscope: the camera pixel size projected in the sample was 80 nm/pixel, and the NA was 1.4. Integration of the PSF over the size of the camera pixels was simulated. The EMCCD camera noise was simulated following the model of Reference (1). The EMCCD camera quantum gain was taken from the instrument datasheet (0.6); thermal noise, and read-out noise were simulated and calibrated following Reference (1).

(5) The 25 SIM images were averaged to generate a wide-field image. The fairSIM reconstruction algorithm was applied to extract a SR-SIM image. The simulated DNA maps (fluorescence intensity signals) were extracted from the horizontal lines where the labels were located in step (3).

#### 3.2 Photo-physical model for simulating other types of super-resolution microscopy

In order to simulate the performance of other types of super-resolution microscopy methods for bacteriophage identification, we modeled the dye behavior as a three-state photo-physical model. The three-state dye model consisted of a bright state, a bleached state, and a dark state. This model allowed us to test two further types of super-resolution microscopy: blinking-based and bleaching-based.

In order to simulate bleaching-based super-resolution microscopy, we used localization by reverse photo-bleaching, reported elsewhere. (2, 3) The average bleaching time was experimentally determined to be 7.5 seconds (i.e. 150 frames, at 50 ms of simulated exposure time). In this method, localization starts at the last frame where almost all dyes are bleached and only isolated emitters are present. These isolated emitters are fitted with a 2D Gaussian using the Localizer software (4) and subtracted from the remaining images. Subsequently the next-to-last image is subjected to the same analysis, and so on until the first image is handled. The localization uncertainty was determined by calculating the standard deviation of the distance between the localized molecules and the DNA fragment. The location of the DNA fragment was determined by fitting a straight line through the dye locations.

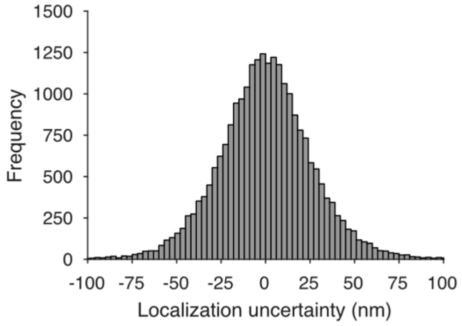


**Figure 10:** Distribution of localizations perpendicular to DNA orientation (in nm).

In order to simulate blinking-based super-resolution microscopy, we set the on-time of the dye to 3 frames and the off-time to 12 frames. We subjected the blinking images to SOFI analysis. (5) Second order SOFI can achieve a similar resolution improvement as SR-SIM, while still being relatively fast (a few 100s of frames are sufficient). Although we found that the simulated performance of using SOFI matched closely to the simulated results for SR-SIM, we could not replicate these results experimentally. In our experiments we noted that the switching of the dyes between bright and dark state was not as predictable as in an ideal three-state system (see Figure 8). Furthermore, this analysis is heavily dependent on the type of dyes and their photo-physical parameters.

#### 3.3 Sensitivity improves with resolution

Figure 11 shows the assignation matrices for simulated data sets using imaging methods with different resolution. For each data set, 1000 DNA fragments were simulated per bacteriophage. The matching significance test was performed with a significance level $\alpha_1 = 0.001$. True positive and false positive matches were counted and are shown as fractions of the number of simulated DNA fragments. Assignation sensitivity clearly improves together with signal resolution (i.e. the on-diagonal elements of the assignation matrices increase)

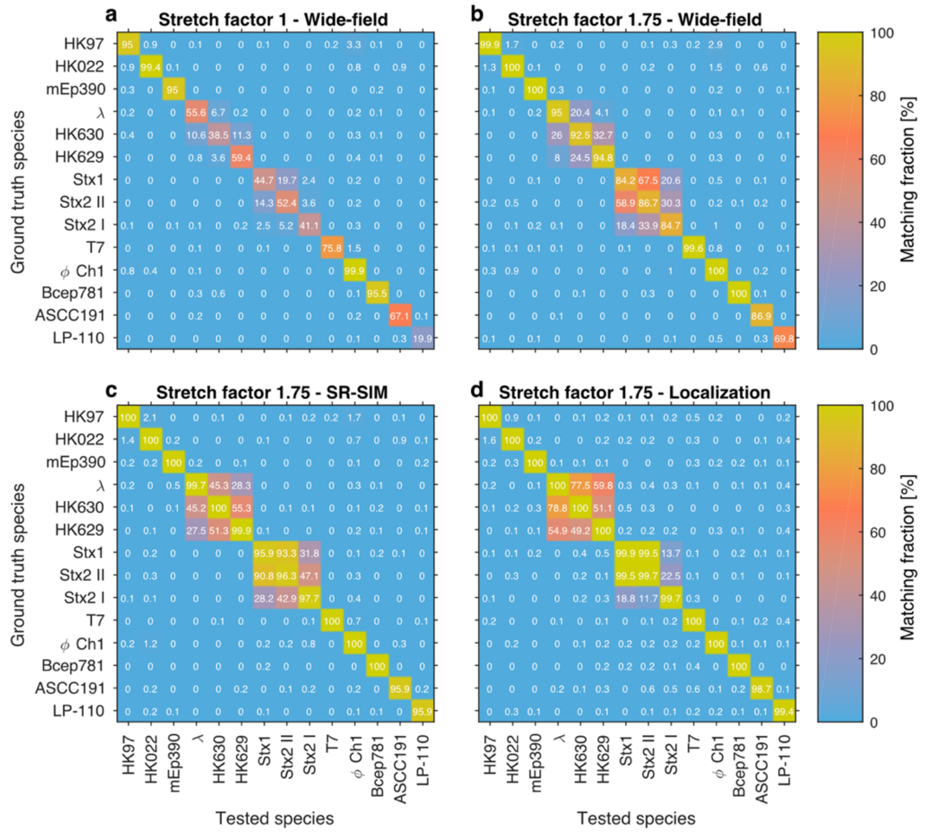


**Figure 11:** Assignation matrices yielded by the matching significance test for different simulated data sets. Significance threshold $\alpha_1 = 0.001$. (a) Unstretched DNA fragments imaged by wide-field microscopy. (b) Overstretched DNA fragments (stretch factor 1.75) imaged by wide-field microscopy. (c) Overstretched DNA fragments (stretch factor 1.75) imaged by SR-SIM microscopy. (d) Overstretched DNA fragments (stretch factor 1.75) imaged by localization microscopy.

#### 3.4 False matching rate increases with resolution for related species (matching significance test only)

In order to investigate the performance of the matching significance test for the identification of closely related species, we simulated data for bacteriophage lambda and for the species listed in Table 2 which are closely related to lambda. Figure 12 shows the false matching rate (for three different significance thresholds) observed when we tried to identify the bacteriophages using only the matching significance test. False matching rate increases when resolution increases for closely related microbial species.

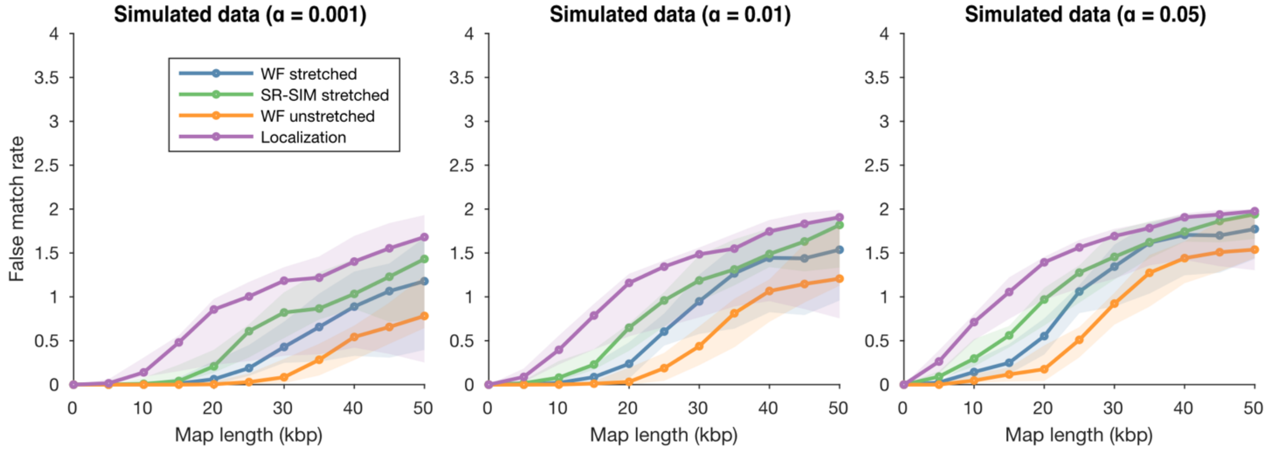


**Figure 12:** False matching rate for closely related species increases with improved resolution when only the matching significance test of the identification method is used at different significance levels (different thresholds for the *p*-values). The shaded regions are delimited by the minimal and maximal false matching rate values for the 3 species concerned.

#### 3.5 Simulated sensitivity as a function of labeling efficiency

Figure 13 shows the effect of the DNA labeling efficiency on the sensitivity of the matching significance test at different significance thresholds. Similar to DNA fragment length, an improved resolution lowers the requirements in terms of labeling efficiency.

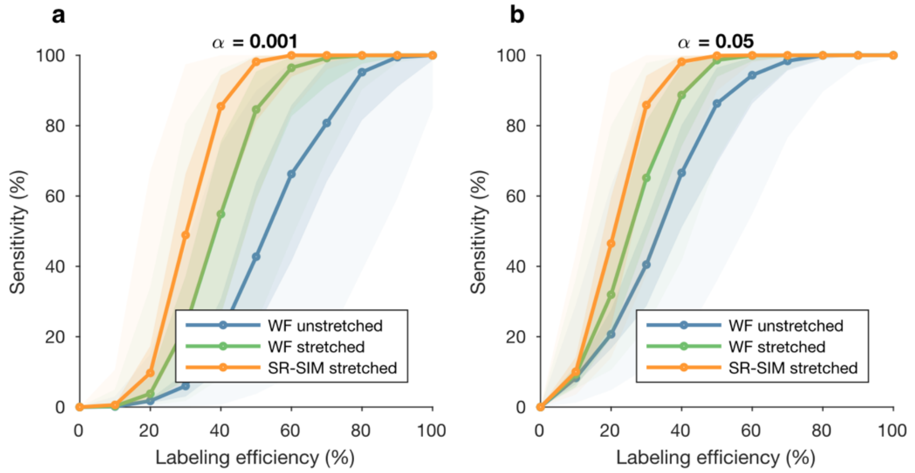


**Figure 13:** Bacteriophage identification sensitivity for 10 species as a function of labeling efficiency (1000 maps per species). (a) Significance threshold $\alpha_1 = 0.001$ (b) Significance threshold $\alpha_1 = 0.05$. The darker shaded regions are circumscribed by the 25-th and 75-th percentile of the sensitivity values obtained for the set of 10 different species. The lighter shaded regions are delimited by the minimal and maximal sensitivity values obtained for the set of 10 different species.

#### 3.6 Specificity increases by resampling the highest significant matching score

Comparing Figure 11 and Figure 14, it can be seen how specificity increases by applying the resampling step of our bacteriophage identification procedure (i.e. the off-diagonal elements of the assignation matrices decrease).

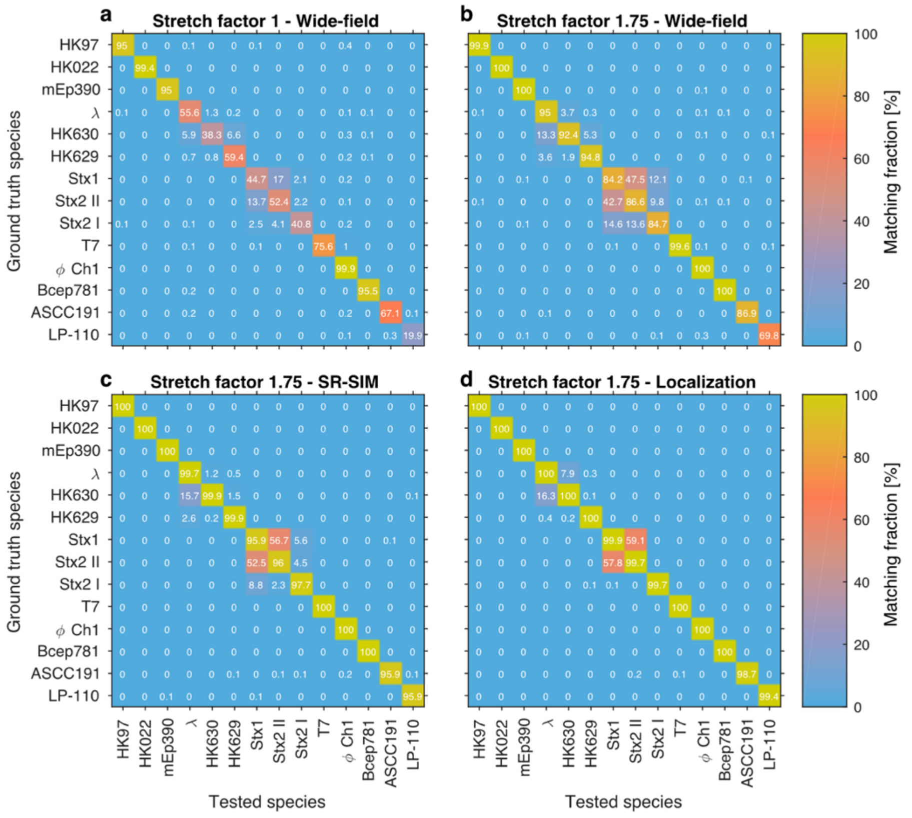


**Figure 14:** Assignation matrices yielded by the matching significance test and the resampling step for different simulated data sets. Significance thresholds $\alpha_1 = \alpha_2 = 0.001$. (a) Unstretched DNA fragments imaged by wide-field microscopy. (b) Overstretched DNA fragments (stretch factor 1.75) imaged by wide-field microscopy. (c) Overstretched DNA fragments (stretch factor 1.75) imaged by SR-SIM microscopy. (d) Overstretched DNA fragments (stretch factor 1.75) imaged by localization microscopy.

#### 3.7 Localization microscopy: dynamic programming versus cross-correlation matching

A frequently used method for matching DNA maps containing localization data is alignment by dynamic programming algorithms. (6, 7) In these cases every gap between two labelled locations is compared to a reference gap, and a score is assigned. These algorithms can accommodate missing labels and false labels by merging two gaps together, as shown in Figure 15.
As an example, Valouev *et al*. published a variation of the Smith-Waterman algorithm (8) which computed the alignment score based on a likelihood ratio for the fragment being the same as in the expected map or random. (7) This model assumes an exponential distribution for the label size and a Poisson process for both the false label and the missing label appearance. The algorithm was designed for restriction mapping. However, the same assumptions can be made for the DNA methyltransferase labeling method used here. We did, however, modify the matching score by adjusting the error in gap sizing (which derives from localization uncertainty in our case), and removed the probability of missing fragments (a specific case for restriction mapping, where small molecules would disassociate from the surface). The resulting equation for the matching score is the following:

$$LR(x, y|m, n) = \frac{\sqrt{\frac{\pi\sigma}{\delta} x^{m-1} e^{\gamma y}}}{(1-\theta)^n \tau^m \frac{x}{\tau} (y\gamma)^{m-1}} \exp \frac{(x-y)^2}{(2\sigma)^2}$$

where $x$ and $y$ refer to the size of the measured map and expected map respectively, $m$ and $n$ refer to the number of false and missing labels. $\theta$ denotes the labeling efficiency, $\gamma$ the false positive rate, $\tau$ the expected size of the gap, $\delta$ the maximal number of consecutive false labels, and $\sigma$ the localization uncertainty.

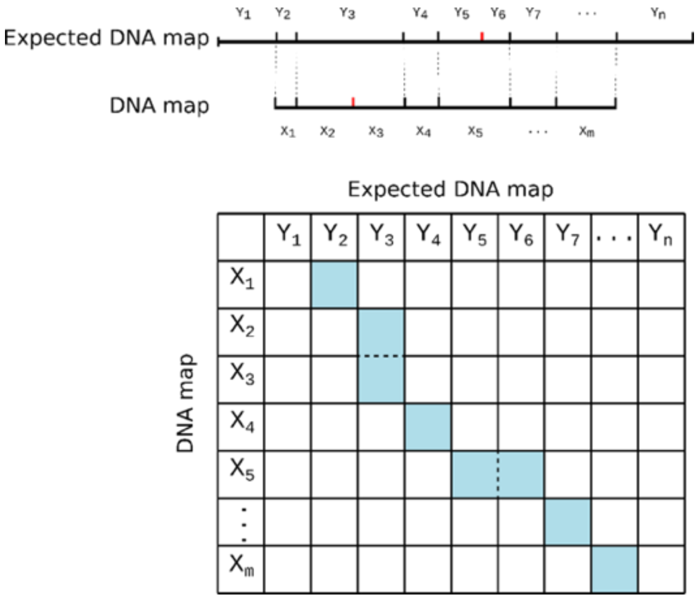


**Figure 15:** Schematic illustration of a dynamic programnming algorithm used for aligning a measured DNA map to an expected DNA map. The false label between gap $X_2$ and $X_3$ and the missing label between gap $Y_5$ and $Y_6$ (highlighted by the dashed lines) can be dealt with by merging the two neighbouring gaps.

Since our approach for calculating a *p*-value for the matching score as well as our method for resampling the highest significant matching score are model- and distribution-free, we can directly apply them to dynamic programming alignment algorithms. In order to assess the performance of dynamic programming alignment for bacteriophage identification, we compared the Smith-Waterman-like algorithm described before to the cross-correlation analysis using a set of simulated localization data. For the cross-correlation analysis, we convolved the localization data with a 2D Gaussian having a FWHM equal to the localization uncertainty (25 nanometers, or 41 basepairs).

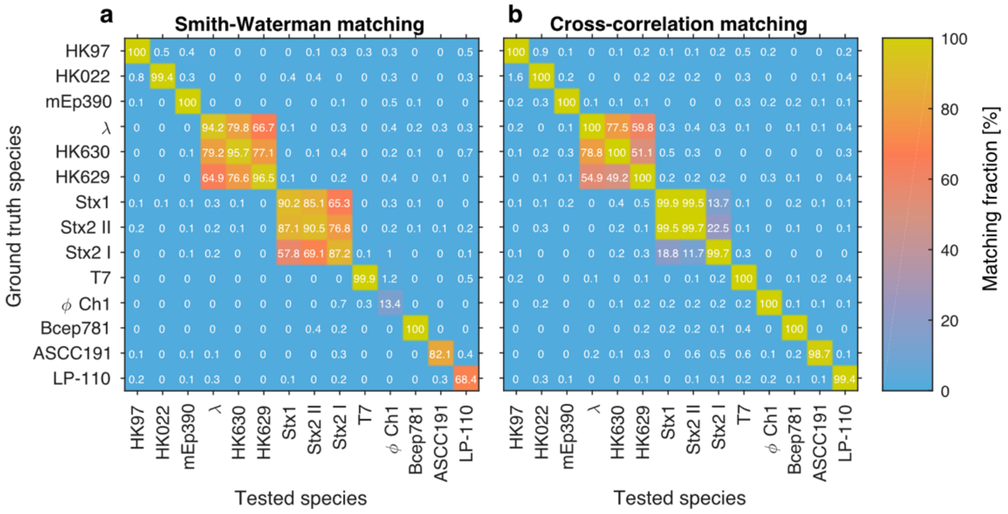


**Figure 16:** Assignation matrices yielded by the matching significance test for localization simulated data sets. Significance threshold $\alpha_1 = 0.001$. (a) DNA fragments imaged by localization microscopy and aligned by dynamic programming using a modified Smith-Waterman function for scoring. (b) DNA fragments imaged by localization microscopy and aligned by cross-correlation matching after Gaussian convolution.

Clearly cross-correlation matching yielded superior results compared to Smith-Waterman-like matching. Most notable is the difference for φCh1, one of the high labeling density species which shows a sensitivity of below 20% when a dynamic programming algorithm is used and a sensitivity of 100% when cross-correlation matching is exploited. Furthermore, this version of the Smith-Waterman alignment requires an estimate of the labeling efficiency and false positive label rate, which further complicates the analysis in real-world scenarios.

#### 3.8 Speed comparison between dynamic programming and cross-correlation

In order to compare the computational time required for DNA matching by both dynamic programming and cross-correlation analysis, we generated random DNA maps of a fixed size with the same labeling density as bacteriophage lambda and ran the alignment 100 times for 10 of such random DNA maps. The computational time was calculated as the average of all 1000 alignments. As shown in Figure 17, the computational time is significantly higher for dynamic programming (almost 2 orders of magnitude longer than for the cross-correlation analysis for a similarly-sized genome). Furthermore, it should be noted that the dynamic programming algorithm scales with the number of labels on the measured DNA map and the expected DNA map, but not with the length (in basepairs) of the two maps. Thus, a high labeling density increases the computational time. The dynamic programming alignment algorithm has a complexity of $O(nm\delta^2)$, where $n$ and $m$ are the number of labels on the measured DNA map and the expected DNA map, respectively, and $\delta$ is the maximal number of consecutive false labels.

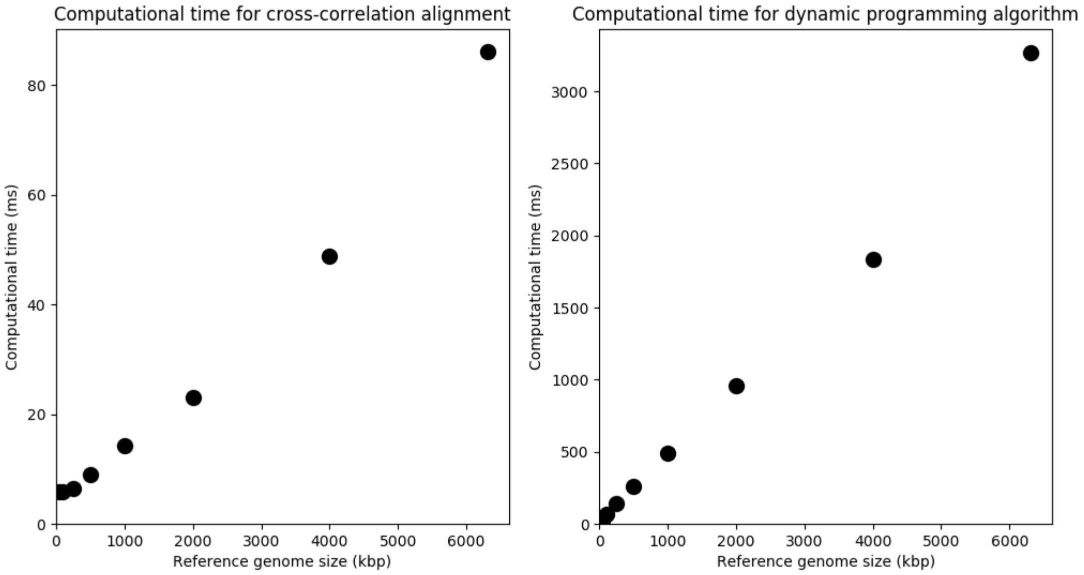


**Figure 17:** Computational time for cross-correlation alignment (left) and dynamic programming alignment (right) as a function of the size of the genome of the reference species.
